# Dynamic single-cell RNA sequencing reveals BCG vaccination curtails SARS-CoV-2 induced disease severity and lung inflammation

**DOI:** 10.1101/2022.03.15.484018

**Authors:** Alok K. Singh, Rulin Wang, Kara A. Lombardo, Monali Praharaj, C. Korin Bullen, Peter Um, Stephanie Davis, Oliver Komm, Peter B. Illei, Alvaro A. Ordonez, Melissa Bahr, Joy Huang, Anuj Gupta, Kevin J. Psoter, Sanjay K. Jain, Trinity J. Bivalacqua, Srinivasan Yegnasubramanian, William R. Bishai

## Abstract

COVID-19 continues to exact a toll on human health despite the availability of several vaccines. Bacillus Calmette Guérin (BCG) has been shown to confer heterologous immune protection against viral infections including COVID-19 and has been proposed as vaccine against SARS-CoV-2 (SCV2). Here we tested intravenous BCG vaccination against COVID-19 using the golden Syrian hamster model together with immune profiling and single cell RNA sequencing (scRNAseq). We observed that BCG reduced both lung SCV2 viral load and bronchopneumonia. This was accompanied by an increase in lung alveolar macrophages, a reversal of SCV2-mediated T cell lymphopenia, and reduced lung granulocytes. Single cell transcriptome profiling showed that BCG uniquely recruits immunoglobulin-producing plasma cells to the lung suggesting accelerated antibody production. BCG vaccination also recruited elevated levels of Th1, Th17, Treg, CTLs, and Tmem cells, and differentially expressed gene (DEG) analysis showed a transcriptional shift away from exhaustion markers and towards antigen presentation and repair. Similarly, BCG enhanced lung recruitment of alveolar macrophages and reduced key interstitial macrophage subsets, with both cell-types also showing reduced IFN-associated gene expression. Our observations indicate that BCG vaccination protects against SCV2 immunopathology by promoting early lung immunoglobulin production and immunotolerizing transcriptional patterns among key myeloid and lymphoid populations.

## Introduction

COVID-19, the current global pandemic caused by the novel coronavirus SARS-CoV-2 (SCV2), has precipitated a severe health and economic crisis worldwide, dramatically affecting the lives of billions of people across all continents. Despite the widespread use of vaccines since December 2020, the limited efficacy of certain licensed vaccines and the development of novel SCV2 variants capable of breakthrough infections in vaccinated individuals poses a continuing public health threat (1, 2). Further, there are indications that SCV2 and future variant coronaviruses may never be fully eliminated, and thus repetitive vaccination campaigns with updated vaccines may be necessary as occurs with influenza (3, 4).

SCV2 infection of ACE2 and TMPRSS2 expressing airway epithelial cells results in pyroptotic cell death and the subsequent release of infectious viral particles, damage-associated molecules (DAMPS), and inflammation inducing IL1-*β* (5, 6). The release of DAMPs and viral particles and their sensing by epithelial cells and alveolar macrophages results in a local exaggerated proinflammatory immune response primarily driven by cytokines (IL-6 and IFN-*γ*) and chemokines (CCL2 and CXCL10) (7). This is followed by infiltration of monocytes, natural killer (NK) cells and lymphocytes from the peripheral blood to infected tissues causing even more pronounced inflammation eventually causing organ damage in individuals with severe disease (7). Longitudinal studies in infected individuals during the active and recovery phases of the disease have revealed that early host immune responses are dominated by cells of innate immune system, including neutrophils, monocytes, plasmacytoid dendritic cells (pDCs), and NK cells (8, 9), while adaptive immune response are important for viral clearance and development of long term T and B cell memory responses (10).

Because BCG has been shown to impart heterologous immunity and to protect against viral infections (11, 12), a number of human clinical trials evaluating BCG for protection against COVID-19 were launched in early 2020. One such study now reports that indeed BCG vaccination reduces the risk of COVID-19 diagnoses by as much as 68% compared to unvaccinated controls (13). BCG is also a well-known vaccine adjuvant and has been shown to boost immunogenicity of protein subunit, DNA, and viral vectored vaccines in pre-clinical models (14–16). Hence further studies of BCG as a COVID-19 vaccine either alone or in combination with the current arsenal of specific anti-COVID-19 vaccines may be warranted.

BCG has been shown to reprogram both myeloid cells and NK cells through processes collectively termed “trained immunity”. BCG is rapidly phagocytosed by macrophages and has been shown to elicit both epigenetic and metabolomic modifications that elevate their immune setpoint upon re-challenge with a heterologous antigens including viruses (17, 18). Upon re-challenge with heterologous antigens, BCG-trained macrophages show elevated cytokine release and demonstrate reprogramming towards M1-like phenotypes. These initial events are associated with heterologous B and T-lymphocyte activation, elevated antibody titers (19) and expansion of unconventional T cells such as innate lymphoid cells (ILCs) and mucosa-associated invariant T (MAIT) cells (20, 21). BCG exposure also leads to expanded “trained” populations of hematopoietic stem cells (HSCs) and multipotent progenitors (MPPs) in the bone marrow that confer enhanced protection against subsequent pathogen challenges (22).

Randomized control studies of BCG vaccination in humans have demonstrated improved control of live attenuated yellow fever virus (23), live-attenuated influenza A (H1N1) (24), and human papilloma virus (25). Furthermore, animal studies have similarly demonstrated a BCG benefit against at least eight viruses including two positive-sense, single-stranded RNA viruses (Cardiovirus A and Japanese encephalitis virus, neither of which is closely related to coronaviruses) (17,26,27). Two recent reports show that subcutaneous BCG does not protect in mouse models of SCV2 infection (28, 29), but that intravenous BCG does confer protection in mice (29). A recent hamster study showed that both subcutaneous and intravenous BCG vaccination were ineffective in SCV2 disease prevention (30). These prior studies did not evaluate the comprehensive cellular immune landscape using robust, global immune profiling transcriptomic tools such as single cell RNA sequencing (28–30).

In this study, we employed the non-lethal, self-resolving golden Syrian hamster model to evaluate the ability of intravenous BCG to protect against SCV2. We evaluated its impact on lung viral titers, tissue pathology using histopathology, immunohistochemistry (IHC) and PET/CT (positron emission tomography/computed tomography), and cell populations by flow cytometry and single-cell RNA sequencing (scRNAseq). Our results reveal that BCG-vaccinated animals had significantly reduced viral loads and tissue pathology, reduced lung T cell lymphopenia, and a significant reprogramming of myeloid cell subsets. Moreover, we observed that during SCV2 infection BCG uniquely recruits immunoglobulin-producing plasma cells to the lung suggesting accelerated antiviral antibody production. BCG vaccination also resulted in elevated levels of several types of lung CD4^+^ T cells (Th1, Th17, Treg, CTLs, and Tmem), and these cells showed modified transcriptional programs towards antigen presentation and away from exhaustion. Similarly, BCG enhanced lung recruitment of alveolar macrophages and reduced key interstitial macrophage subsets with both cells also showing reduced IFN-associated gene expression. Our results indicate that BCG vaccination has a tolerizing effect on SCV2-mediated lung inflammation and that it may accelerate antiviral antibody responses.

## Results

### BCG vaccination reduces viral load and lung pathology and alters lung immune cell infiltration in SCV2-infected hamsters

We set out to determine whether prior BCG vaccination alters the course of SCV2 infection in the golden Syrian hamster model (**Fig. 1A**) which has previously been shown to exhibit a self-resolving, and non-lethal form of COVID-19. We evaluated lung histopathologic changes at D4 and D7 post SCV2 challenge with and without prior BCG vaccination. As shown in **Fig. 1B-C**, bronchopneumonia was present in 100% of unvaccinated hamsters on both D4 and D7 but was absent in all BCG-vaccinated hamsters on D4 and found in only 25% on D7. Similarly, 100% of unvaccinated hamsters showed neutrophilic infiltration at D4 while only 50% of BCG-vaccinated hamsters showed this finding (**Fig. 1D**). In contrast BCG vaccination showed a trend for increased macrophage infiltration (80% at D4) while only 60% of unvaccinated animals showed this finding (**Fig. 1E**). Regarding the well-described SCV2-mediated lymphopenia, BCG vaccination was able to partially correct infiltrating lung lymphocytes with BCG-vaccinated hamster lungs also showing significantly higher levels of lung lymphocytes (CD3 positivity) both by traditional histology and immunohistochemistry (**Fig. 1F, Fig. S1A**), and this was true for perivascular lymphocytic infiltration as well (**Fig. S1B**). As expected, all BCG vaccinated hamsters showed granuloma formation, a phenomenon that was uniformly absent in unvaccinated animals (**Fig. S1C)**. Thus, prior BCG vaccination reduced SCV2-induced bronchopneumonia with its concomitant neutrophilic infiltration while simultaneously preventing lung lymphopenia and enhancing lung macrophage levels.

**Figure 1.**
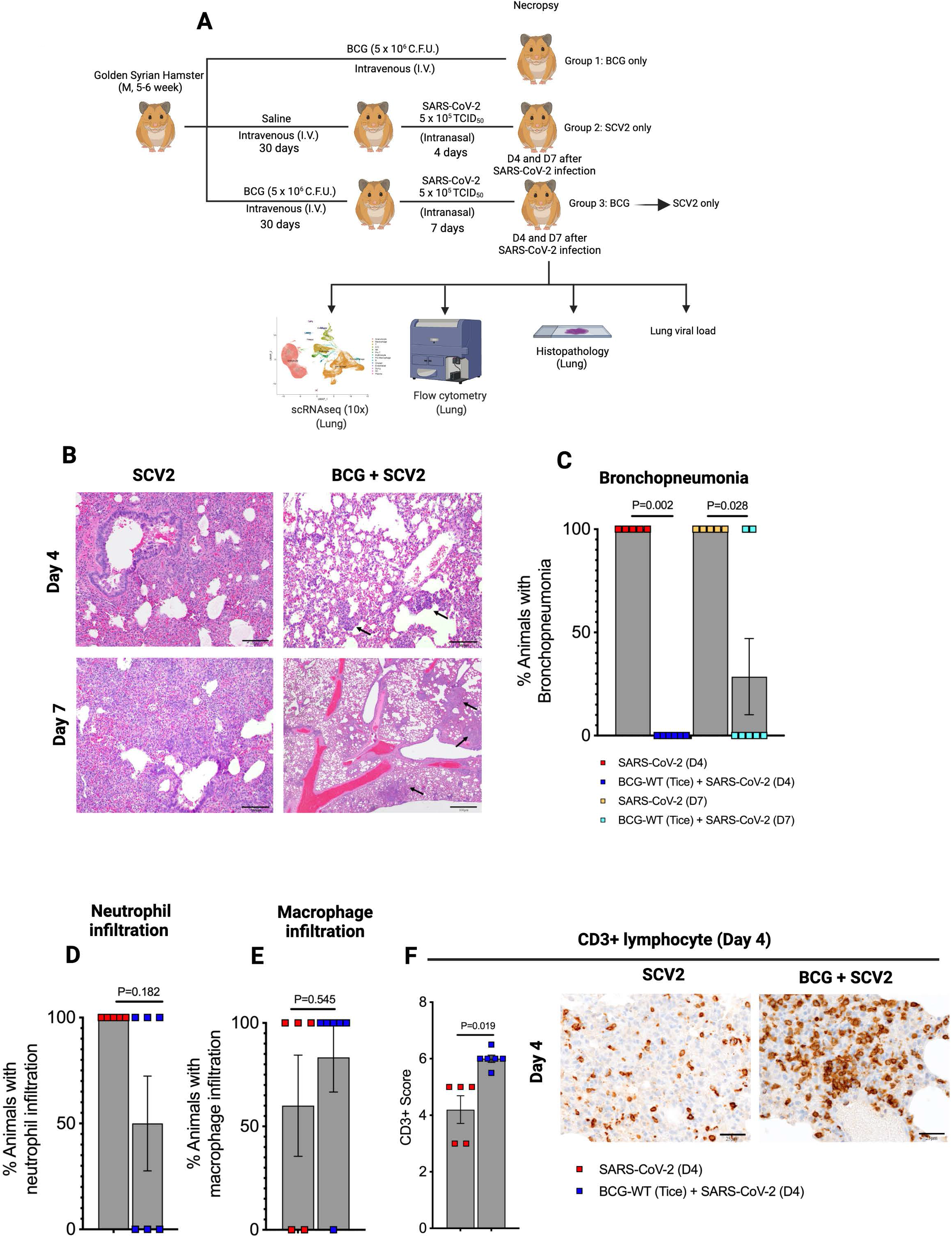
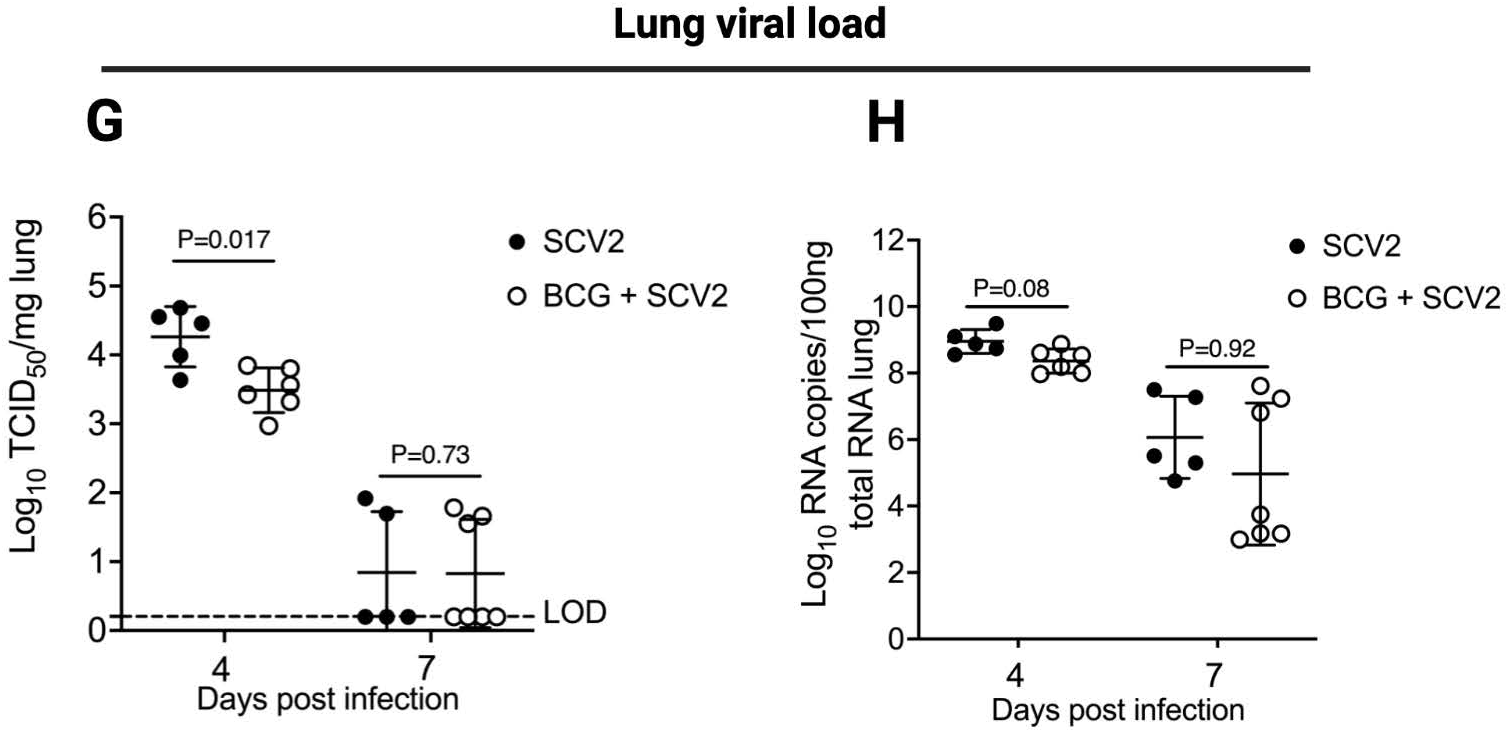
BCG vaccination diminishes SARS-CoV-2 (SCV2) infection induced lung inflammation and viral load in golden Syrian hamsters. **A.** Experimental layout. **B-C.** Bronchopneumonia at day 4 (D4) and day 7 (D7) after SARS-CoV-2 infection, mild local inflammation (arrows) in BCG + SCV2 lungs. Neutrophils (**D**) and macrophage (**E**) infiltration at D4 after SARS-CoV-2 infection. **F**. CD3^+^ lymphocyte positivity scores at D4 in hamster lungs after SARS-CoV-2 infection. **G-H.** Bar graph showing SARS-CoV-2 viral load in the lung at D4 and D7 post infection in non-vaccinated and BCG-vaccinated hamsters. Briefly, Infective viral particles (**G**) were quantified from a standard of infectious virus and expressed at TCID_50_ equivalents per mg lung tissues. Dotted lines indicate lower limit of detection (LOD). Viral RNA levels (**H**) were determined in the lungs, normalized against human RNaseP and were transformed to estimate viral RNA copies per 100 ng of total lung RNA. Data points represent number of animals used per group. Statistical analysis was done using two-sided Fisher exact test (**C-E**), Welch’s t-test (**F**) or two-tailed Student’s t-test (**G-H**) (significant when P < 0.05). Percent animals and 95% confidence intervals (CI) showing pathology were (C) bronchopneumonia in the BCG-SCV2 group at D4 0% (95% CI: 0, 45.9%), at D7 28.6% (95% CI: 36.7, 71.0%); (**D**) neutrophil infiltration in the BCG-SCV2 group at D4 50% (95% CI: 11.8, 88.2%); (E) macrophage infiltration in the BCG-SCV2 group 83.3% (95% CI: 35.9, 99.6%) and in the SCV2 only group at D4 60.0% (95% CI: 14.7, 94.7%). A schematic of CD3 positivity (**F**) is given in methods section.

We also evaluated the effect of BCG vaccination on lung viral load by measuring infectious viral particles (TCID_50_) and viral RNA in lung homogenates. We observed a significant decrease in TCID_50_ in BCG vaccinated animals at D4 (**Fig. 1G**), and a trend of reduced viral mRNA (**Fig. 1H**) at this same time point. TCID_50_ and viral mRNA levels were significantly lower at D7 compared to D4 and levels BCG-vaccinated animals did not differ from the unvaccinated ones likely because of progressive disease resolution in both groups.

### BCG vaccination blunts SCV2-mediated lung T cell lymphopenia, enhances macrophage lung recruitment, and reduces lung infiltration by granulocytes

To assess overall cell population fluxes in hamster lungs, we performed multicolor flow cytometry. We found that lung T cell infiltration (CD3^+^ cells) was dramatically boosted by BCG vaccination in the absence of SCV2 (45% of live cells) compared to age-matched healthy animals (18%) (**Fig. 2A**). In unvaccinated animals challenged with SCV2, the total lung CD3^+^ population was reduced at D4 and D7 to 4-6%; while BCG vaccinated, SCV2 challenged animals showed a trend to have nearly twice as many CD3^+^ cells (10-12%) at these same time points (**Fig 2A**) supporting our histopathological observations. This same effect of BCG preventing T cell depletion in the lung was present to an even greater extent among lung CD4^+^ T cells (**Fig. 2B**). We did not observe this same pattern in hamster spleens suggesting that the observed effects of BCG vaccination were limited to the site of active infection in the lungs (**Fig. S2 A-B**). Similarly, BCG vaccination in the absence of SCV2 infection prompted a pronounced macrophage recruitment to the lung (55% of live cells) compared to healthy (unvaccinated) animals (5%) (**Fig. 2C**). And while SCV2 infection led to a modest macrophage lung recruitment by D7 (12% of live cells), previously BCG vaccinated animals showed significantly elevated macrophage populations in the lungs (30% of live cells) on both D4 and D7 (**Fig. 2C**). In contrast, for granulocytes, we found that SCV2-challenged, unvaccinated hamster lungs revealed dramatic infiltration by polymorphonuclear leukocytes (PMNs) on D4 (80% of live cells) and D7 (72%) (**Fig. 2D**) in accordance with the profound bronchopneumonia observed (**Fig. 1B-C**). Pre-vaccination with BCG had the effect of limiting the SCV2 mediated granulocytic infiltration to levels of 30% on D4 and 32% on D7 which were levels essentially equivalent to those in the lungs of vaccinated, unchallenged hamsters. A modest increase in macrophages at both D4 and D7 in the splenic compartment, not statistically significant, was observed in BCG-vaccinated, SCV2-infected animals compared to unvaccinated SCV2 infected animals (**Fig. S2C**). Splenic granulocytes trended to be lower in BCG vaccinated animals compared to unvaccinated animals (**Fig. S2D)**, similar to the observations in the lungs.

**Figure 2.**
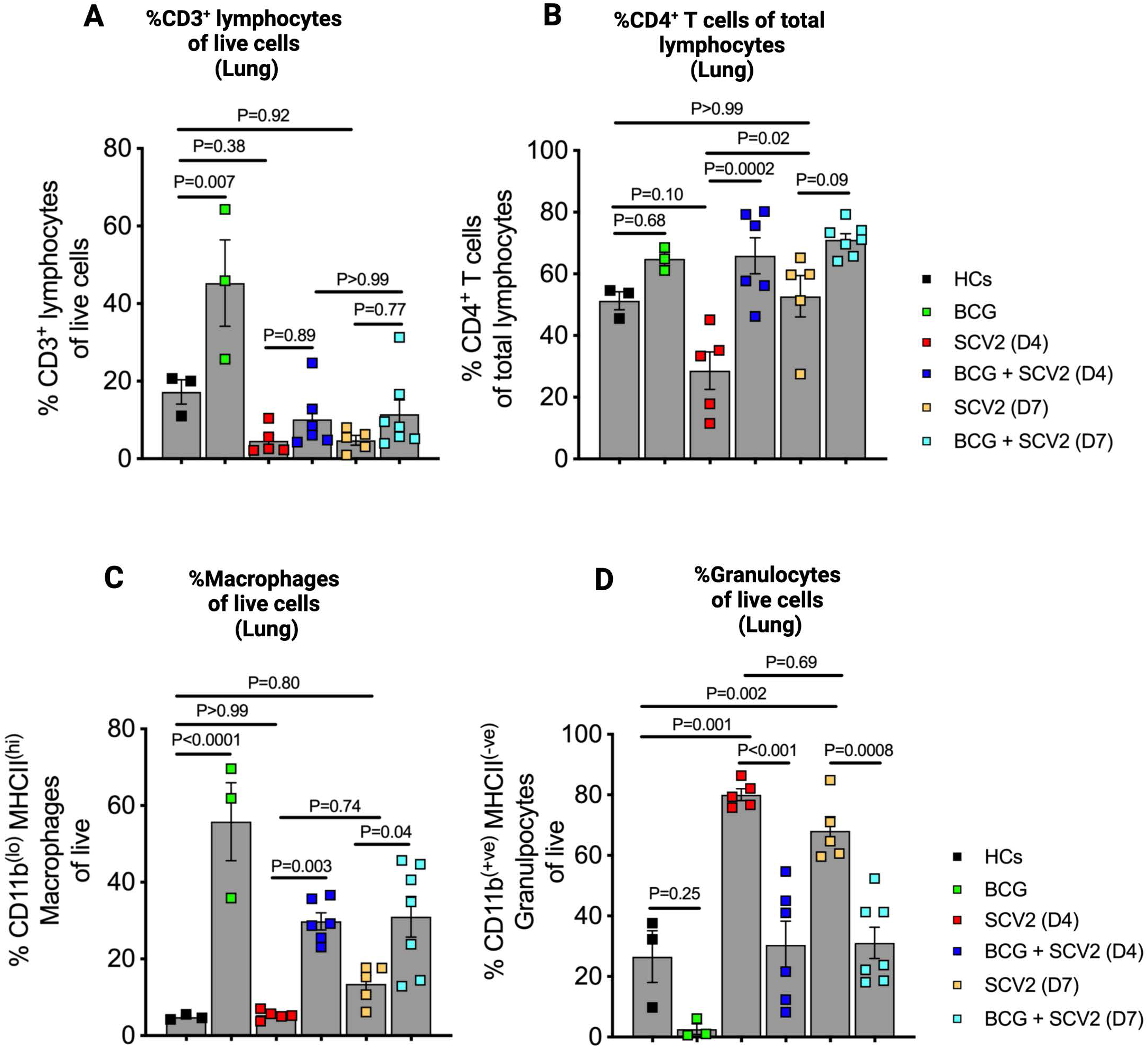
Differential abundance of lymphocytes, macrophages and granulocytes in BCG vaccinated golden Syrian hamster lungs after SARS-CoV-2 infection. Differential percentages of **A.** CD3^+^ lymphocytes. **B.** CD4^+^ T cells within the lymphocyte compartment. **C.** macrophages (CD11b^(lo)^ MHCII^(hi)^ of live cells) and, **D.** granulocytes (CD11b^(+ve)^ MHCII^(-ve)^ of live cells) in the lung of hamsters. Single cells preparations from lung tissues following necropsy were investigated for different lymphoid and myeloid cells using multicolor flow-cytometry. The data are presented as mean values ± S.E.M. Statistical analyses were done using one-way ANOVA with the p-values indicated.

### Single cell transcriptional profiling in lung cells in BCG vaccinated and unvaccinated hamsters

To obtain a higher resolution of pulmonary immune responses linked to BCG immunization during SCV2 infection, hamster lungs were examined using droplet-based single cell RNA sequencing (10X Genomics) during the peak (day 4) and resolution (day 7) phases of COVID-19. BCG-vaccinated, unchallenged hamster lungs were also studied. A total of 13 hamsters were evaluated in the following three groups: BCG vaccination only (3 animals), SCV2 infection only (2 animals on D4; 2 on D7), and BCG vaccination with SCV2 challenge (3 animals on D4; 3 on D7). Sequencing was performed in a total of 194,536 cells, and after filtering out low-quality cells, red blood cells, and doublets 112,928 cells were analyzed. (**Table S1**). The mean number of cells analyzed per animal in BCG vaccinated lungs, SCV2 infected lungs and BCG-vaccinated + SCV2 infected lungs was 11,249, 8,643, and 7,434 respectively.

### BCG reverses lymphopenia, macrophage depletion, and loss of structural cell types due to SCV2 infection

By graph-based clustering of uniform manifold approximation and projection (UMAP), transcriptomes of 17 major cell types or subtypes were identified in hamster lungs (**Table S2**). These included lymphoid, myeloid, and non-immune cell types (**Fig 3A-C**). These cell clusters were relatively homogenously distributed across all animal lungs at both timepoints and were based on the expression of well-defined canonical genes as shown in **Fig. 3D**. The 5 most highly expressed genes in each cluster are shown in **Fig. S3**.

**Figure 3.**
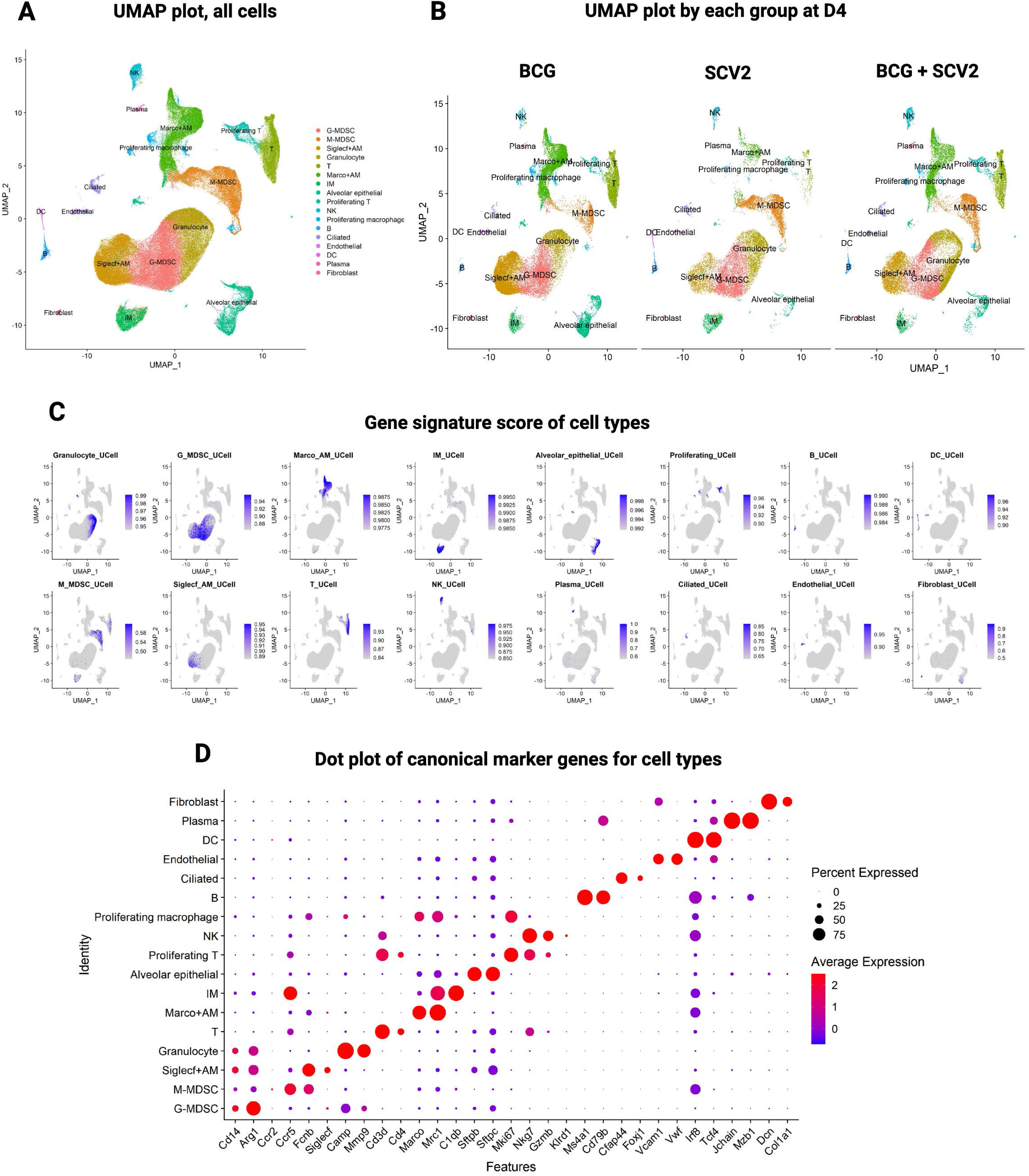
Cellular landscape of SARS-CoV-2 infected hamster lungs after BCG vaccination identified by scRNAseq. **A.** Uniform Manifold Approximation and Projection (UMAP) plot showing identification of 17 different major cell-type in hamster lungs integrated by all samples. **B.** UMAP plot showing lung cellular dynamics in hamster lungs in BCG vaccinated, SARS-CoV-2 (SCV2) infected, and BCG + SARS-CoV-2 (SCV2) infected animals at D4 after SCV2 infection. **C.** UCell score distribution in UMAP plot for cell type marker gene signatures (**listed in Table S2**) evaluated using UCell. **D.** Dot plots of log-normalized expression and fraction of cells expressing selected canonical marker genes for identification of 17 different cell types in hamster lungs across all samples.

### BCG expands Th1, Th17, Treg, CTLs, Tmem, and plasma cell populations during SCV2-infection

Next, we performed subset analyses to evaluate lung immune cells during SCV2 infection in vaccinated and unvaccinated hamsters. We were able to identify 10 distinct lymphocyte cell types (**Fig. 4A, S4A-B**). Among the CD4^+^ T cells in hamster lungs, we were able to distinguish Th1, Th17, Treg (regulatory T cells) and Tmem (memory T cells) cells primarily based on *Ifng*, *Itgb7*, *Foxp3*, and *Lef1* expression, respectively (**Fig. S4C-D**). CTLs (cytotoxic T cells) showed high *Gzmk* and *Gzma* but low *Cd4* expression. Plasma cells were identified by their elevated expression of *Jchain* and *Mzb1* (**Fig. S4C-D**). As may be seen in **Fig. 4A**, both at D4 and D7 post-SCV2 challenge, hamsters that were BCG-vaccinated maintained high levels of Th1, Th17, Treg, CTLs, Tmem, and plasma cells in their lungs. With the exception of plasma cells, vaccination with BCG only (no SCV2 challenge) stimulated high levels of these cell types in the lungs, and these relatively high levels were maintained in hamsters challenged with SCV2. In contrast, a plasma cell response was not observed in BCG-vaccinated, unchallenged animals; rather elevated plasma cell numbers in the lungs were uniquely observed only in BCG-vaccinated, SCV2 infected animals.

**Figure 4.**
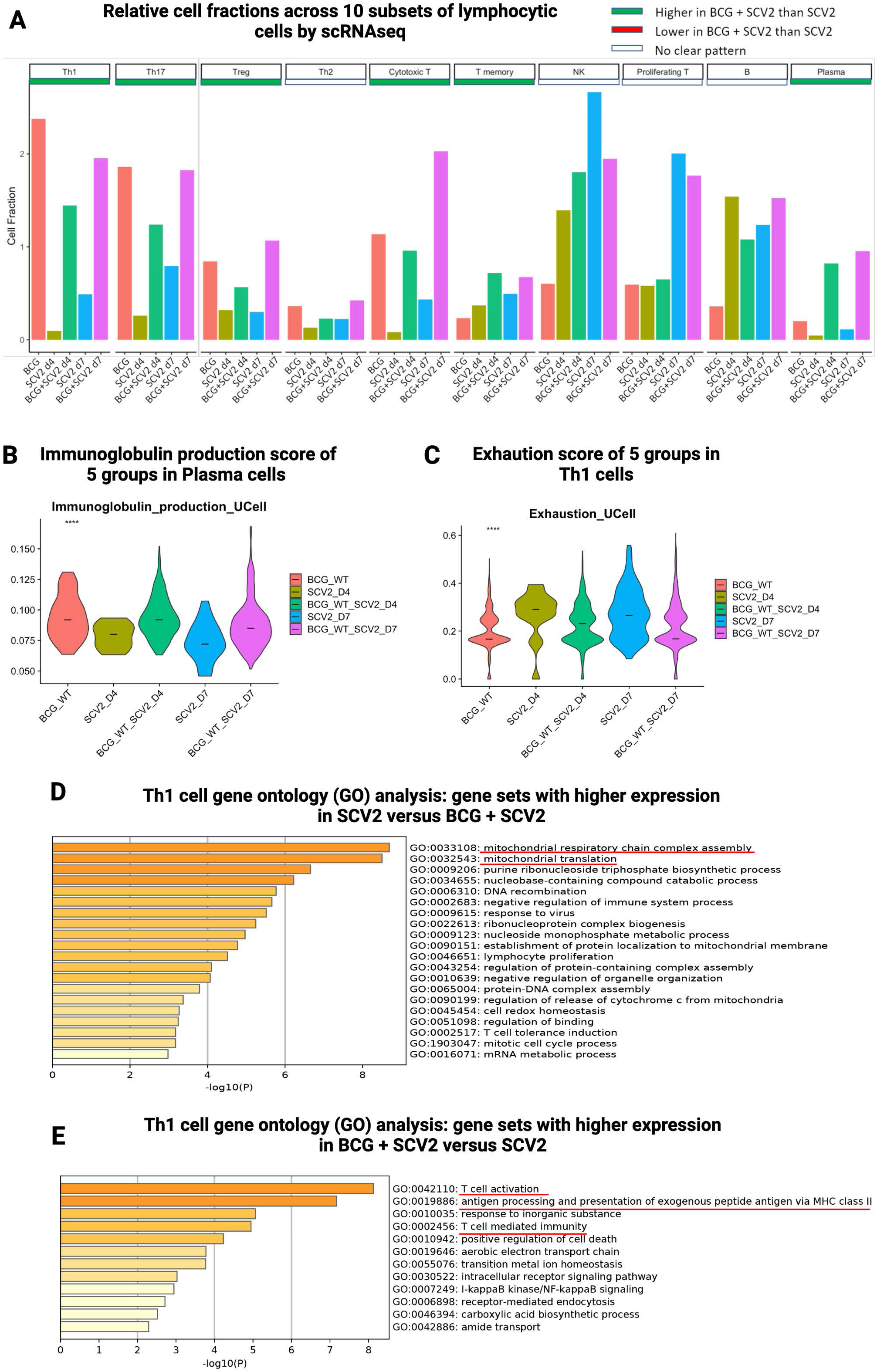
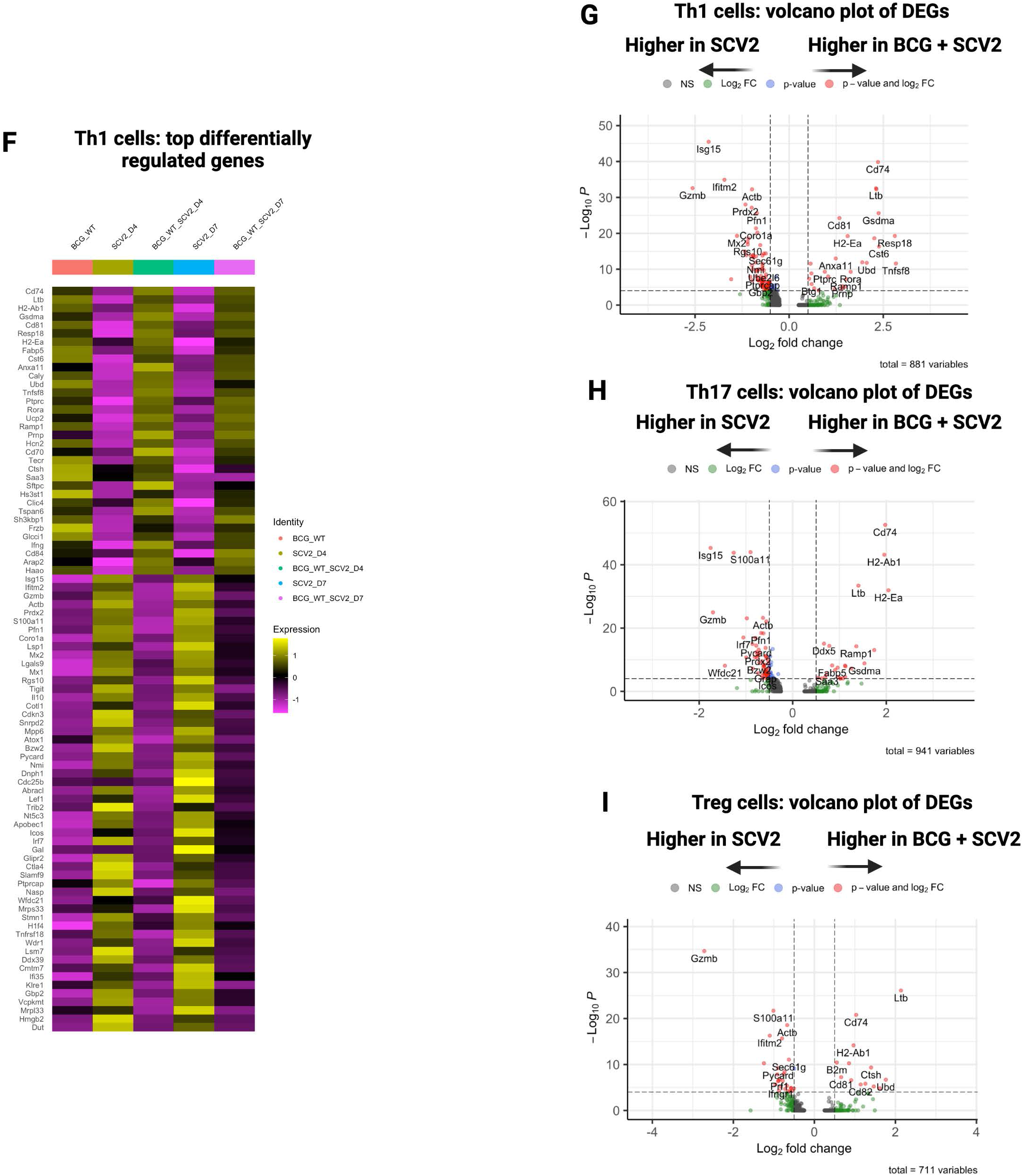
BCG vaccination strongly affects lung lymphoid dynamics and gene activation programs in different lymphoid subsets in SARS-CoV-2 infected hamster lungs. **A.** Bar graph showing average proportion of different lymphocytic cells across groups. Violin plots showing UCell scores for (**B**) immunoglobulin production on plasma cells, and (**C**) exhaustion scores on Th1 cells across different groups. Horizontal lines represent mean values. All differences with P < 0.01 are indicated. **P < 0.01, ***P < 0.001, ****P < 0.0001 (one-way ANOVA). Th1 lymphocyte-linked gene ontology enrichment analysis of biological processes showing gene sets that enriched in (**D**) SARS-CoV-2 (SCV2) versus BCG + SARS-CoV-2 (SCV2), and in (**E**) BCG + SARS-CoV-2 (SCV2) versus SARS-CoV-2 (SCV2) across both time-points. A darker color indicates a smaller P value. F. Heat map depicting the average expression value of top differentially expressed genes (DEGs) across different treatment conditions in Th1 cells. Top DEGs were selected with adjusted P-value < 0.05 and average fold change >0.7. Volcano plots depicting DEGs in (**G**) Th1, (**H**) Th17, and (**I**) regulatory T cells (Tregs). The x-axis indicates log_2_FC of gene expression in BCG + SCV2 compared to SCV2 group.

### Elevation of plasma cell abundance in BCG-vaccinated, SCV2 challenged hamsters

In order to estimate the numbers of lymphocytes in healthy control hamsters, we were able to compare our scRNAseq cell abundance data with previously published data which included healthy hamster controls (31) (**Fig. S5A**). Both Th1 and Treg lymphocytes were scarce in the lungs of healthy hamsters but were strongly recruited to the lung by BCG vaccination both in the presence and absence of SCV2 challenge (**Fig. S5B**). Plasma cells were not seen abundantly in either healthy hamster lungs or unchallenged, BCG-vaccinated hamsters. Thus, an enhanced plasma cell response occurred only in the presence of prior BCG-vaccination followed by SCV2 challenge (**Fig. 4A**). Moreover, an evaluation of gene signatures revealed elevated expression of immunoglobulin production by plasma cells in BCG-vaccinated and SCV2 challenged animals strongly suggesting that these plasma cells appear functionally active in antibody production (**Fig. 4B**).

### Lung CD4^+^ Th1 cells showed reduced exhaustion and mitochondrial stress gene signatures with BCG vaccination

By gene signature and gene ontology (GO) analysis, we found that Th1 cells in unvaccinated, SCV2-challenged showed evidence exhaustion markers (**Fig 4C**) and a high degree of mitochondrial respiratory gene expression and translation (**Fig 4D**), consistent with mitochondrial dysfunction in T cells that have been previously reported (32). In contrast, in Th1 cells from BCG-vaccinated, SCV2 challenged animals, pathways related to T cell activation, T cell immunity, and antigen processing were upregulated suggesting greater functional immunity (**Fig. 4E**).

### Differentially expressed genes (DEGs) in lung CD4 Th1 cells

When we considered specific DEGs in Th1 cells in BCG-vaccinated, SCV2-challenged hamsters (**Fig 4F,G**), we found strong upregulation of genes associated with protection from injury. These include *Cd74,* a gene recently associated with antiviral responses and protection from lung injury (33, 34), *Cd70* a gene associated with T cell activation, *Ltb* (lymphotoxin-beta) a gene known to play a protective role in other viral infection, *Gsdma* (gasdermin A) which has been found to be deficient in autopsy studies of humans with severe COVID-19 (35), and *Ifng* (IFN-*γ*) a prominent Th1 activator. Several of these same genes including *Cd74*, *Ltb*, and *Gsdma* were also strongly expressed in Th17 and Treg lung lymphocytes from BCG-vaccinated, SCV2-challenge hamsters (**Fig. 4H-I**).

In contrast, in Th1 cells from non-vaccinated, SCV2-challenged hamsters, we found DEGs that were associated with markers of exhaustion and ongoing viral infection (**Fig. 4F,G**). Markers of exhaustion included *Cd278* (*Icos*), a gene known to be up-regulated in SCV2 (36), *Ctla4*, a marker found in human SCV2 convalescent plasma (37), and *Tnfrsf18* (GITR). Genes associated with ongoing viral infection included *Ifitm2* (IFN-induced transmembrane protein 2) which has recently been shown to be hijacked by SCV2 and to promote ongoing infection response (38), *Bzw2* (Basic Leucine Zipper and W2 Domain-Containing Protein 2) (39) which is a viral restriction factor shown to interact with the SCV2 M protein, and IFN-responsive regulatory proteins *Mx1* (an IFN induced GTP-binding protein), *Rgs10* (Regulator of G protein signaling-10), and *Irf7* which is known to be involved in transcription activation of viral-inducible cellular genes and is a gene for which polymorphisms known to be associated with severe SCV2 (40).

### Myeloid cell changes mediated by BCG vaccination: enhanced AM abundance and diminished IM lung recruitment

Our scRNAseq analysis of myeloid cells revealed 10 distinct myeloid cell subsets (**Fig. 5A, Fig. S6A-B**). Alveolar macrophages (AMs) are relatively immunotolerant cells that scavenge debris and play an important role in pathogen clearance (41). We identified two clusters of AMs, the first identified by the classic AM marker *Siglecf* (Siglecf+ AMs) and the second identified by the presence of the scavenger receptor *Marco* (Marco+ AMs). BCG vaccination has been shown to induce recruitment of Siglecf+ AMs to the lung in healthy human volunteers (42), and consistent with this, while we observed virtually no Siglecf+ AMs in healthy hamster lungs, this population was strongly induced by BCG vaccination (**Fig. S5C**). In contrast, scavenging Marco+ AMs were present in high numbers in healthy hamster lungs, and BCG vaccination caused ∼2-fold increased in this population (**Fig. S5C**). Unvaccinated, SCV-infected hamsters showed a high degree of depletion of both categories of AMs, but prior BCG vaccination served to prevent the loss of both AMs from the lung (**Fig 5A, S5C**).

**Figure 5.**
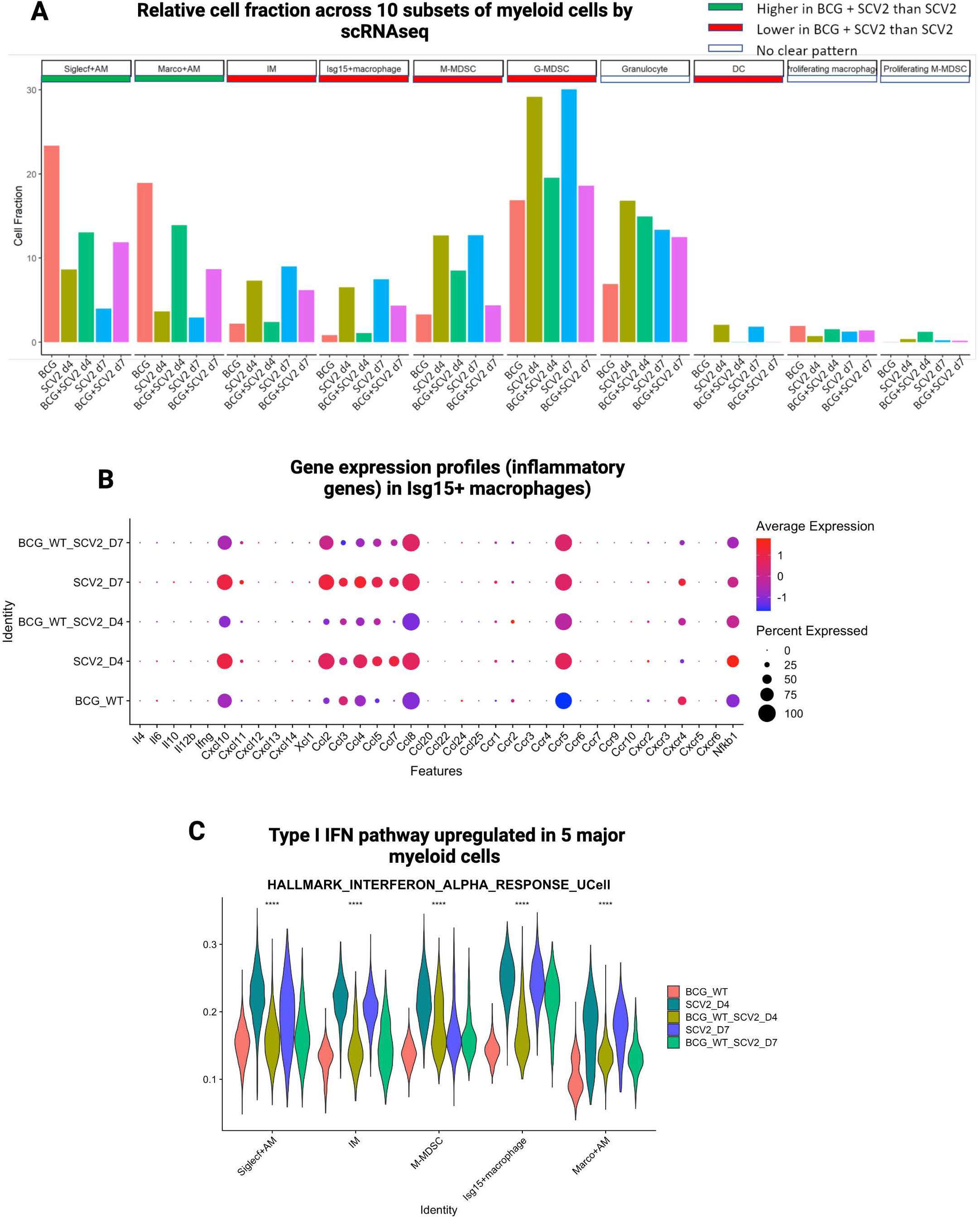
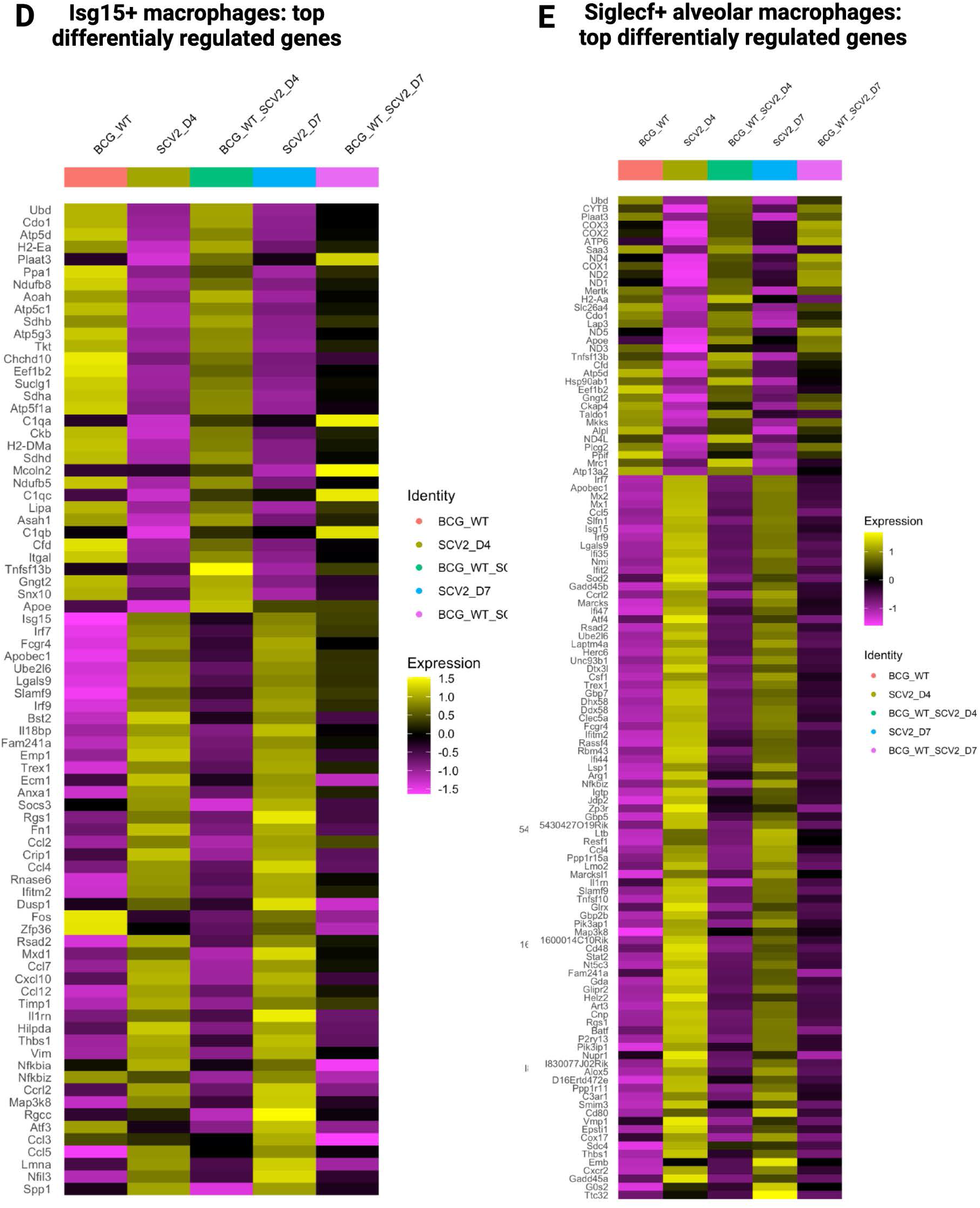
BCG vaccination curtails SCV2 mediated lung inflammation by modulating myeloid cell abundance and phenotypes. **A.** Bar diagram showing average proportion of different myeloid cells across groups. **B.** Dot plot showing the expression levels of a group of chemokines and cytokines in Isg15+ macrophages across different groups. **C.** Violin plots showing UCell gene signature scores of hallmark type I IFN (Interferon-*α*) responses in 5 major myeloid cells in hamster lungs across different groups. All differences with P < 0.01 are indicated. **P < 0.01, ***P < 0.001, ****P < 0.0001 (one-way ANOVA). Heat map depicting the average expression value of top differentially expressed genes (DEGs) across different groups in **(D)** Isg15+ macrophages, and **(E)** Siglecf+ alveolar macrophages (AMs). Top DEGs were selected with adjusted P-value < 0.05 and average fold change > 0.7.

Interstitial macrophages (IM) are more inflammatory than AMs and are recruited from the periphery during lung infection (43, 44). In addition to traditional IMs, we identified a second population which strongly expressed *Isg15* (Isg15+ macrophages) indicating that they are highly responsive to IFN I and hence likely to be hyperinflammatory. Both IMs and Isg15+ macrophages were virtually absent in healthy hamster lungs (**Fig. S5C**) and in unvaccinated hamsters, but both of these cell types were recruited to the lungs in high numbers following SCV2 challenge. Interestingly, prior BCG-vaccination significantly blunted this recruitment (**Fig 5A, S5C**). In keeping with the inflammatory nature of Isg15+ macrophages, we observed that SCV2 infection caused a strong upregulation of *Ccr5*, *Ccl8*, *Ccl7*, *Ccl5*, *Ccl4*, *Ccl2* and *Cxcl10* in IFN-I-responsive Isg15+ macrophages (**Fig. 5B**). These inflammatory chemokines remained at basal expression levels in either BCG vaccinated animals or those challenged with SCV2 after vaccination. These observations support the notion that BCG vaccination leads to an elevated setpoint of scavenging AM in the lungs, but during SCV2 infection it prevents excess recruitment of pro-inflammatory Isg15+ macrophages and IMs which may be involved in SCV2-mediated, lung immunopathology.

### Dendritic cells (DCs)

We observed an increase in dendritic cells following SCV2 infection at both time points. In contrast, levels of DCs remained essentially undetectable with BCG vaccination alone or BCG-vaccination in the presence of SCV2 challenge (**Fig 5A**). Patients with severe COVID-19 demonstrate reduced DC levels in blood possibly because of recruitment to infection sites (45). DCs, especially pDCs, are considered a chief source of IFN I, a cytokine which is found at unusually low levels during human COVID-19 suggesting that DCs have an exhaustion phenotype during SCV2 infection (46, 47).

### Myeloid-derived suppressor cells (MDSCs)

Expansion of MDSCs have been described in humans with severe COVID-19, a phenomenon that correlates with lymphopenia and enhanced arginase activity, while milder forms of disease display a reduced MDSCs count (48, 49). Increased recruitment of both G-MDSCs and M-MDSCs subtypes of myeloid cells are reported in COVID-19 patients and may contribute to acute respiratory distress syndrome (ARDS) (50, 51). BCG vaccination led to reductions of in both M-MDSCs and G-MDSCs among SCV2-challenged hamsters suggesting a protective effect (**Fig. 5A**). The majority of G-MDSCs identified in our data showed marked elevation of *Il1b* and *Arg1* both strong biomarkers of COVID-19 severity (51) (**Figure S3**).

### Myeloid cell DEGs reveal divergent immune programs

Next, we evaluated DEGs in Siglecf+ AM, Marco+ AM, IMs, Isg15+ macrophages, and M-MDSCs. In unvaccinated animals infected with SCV2, there was high transcription of IFN-*α* (IFN-I) response genes (**Fig 5C**); these included *Isg15*, *Irf7*, and *Mx1* as well as chemokine-encoding genes such as *Ccl2*, *Ccl12*, and *Ccl4* and those associated with lymphocyte activation (*Slamf9*) all consistent with ongoing inflammatory responses (**Fig. 5C-D**). However, in vaccinated animals challenged with SCV2, the predominant DEGs were shifted towards those involved in metabolic and repair processes such as *Ubd* (encoding ubiquitin), *Ppa1* (encoding pyrophosphatase 1), *Cdo* (encoding cysteine dioxygenase) or genes for complement factor expression (*C1qa*, *C1qc*) as might be expected in a less inflammatory, repair-oriented environment (**Fig. 5C-D**). Focused analysis of IFN-*α* and IFN-*γ* response gene signature scores revealed that for major myeloid cell subsets, BCG-vaccinated animals showed reduced expression of these interferons (**Fig. 5C**), including for Isg15+ macrophages which had the highest levels of IFN-*γ* expression.

### DEGs in Isg15+ macrophages

In unvaccinated, SCV2-challenged Isg15+ macrophages we observed robust upregulation of type I IFN-dependent ISGs (interferon-stimulated genes), (*Bst2, Mxd1* and *Ifitm2*), ISGs known to be activated following viral genome sensing (*Irf7*, *Fcgr4*, *Slamf9*, and *Irf9),* antiviral restriction factors (*Apobec1),* and inflammatory mediators known to drive COVID-19 severity (*Ccl2*, *Ccl4*, *Ccl7*, *Ccrl*2, *Cxcl10*, and *Ccl12*) (**Fig. 5D**). In contrast, Isg15+ macrophages among BCG-vaccinated, SCV2-challenged hamsters expressed high levels of oxidative phosphorylation genes including *Atp5*, *Atp5c1*, *Atp5f1a* and *Atp5g3* as well as mitochondrial *Chchd10* (a gene predicted to be involved in mitochondrial oxidative phosphorylation) suggesting upregulated oxidative phosphorylation, a characteristic hallmark of BCG training (52). Isg15+ macrophages among BCG-vaccinated animals showed increased expression of mitochondrial genes *Suclg1* (succinyl-CoA ligase), and *Sdhd* (succinate dehydrogenase) suggested increased generation of succinate and fumarate respectively. Both metabolites play extensive roles in epigenetic reprogramming of macrophages, a characteristic also associated with macrophage training. Interestingly, increased expression of *Aoah* (encoding acyloxyacyl hydrolase), a gene usually upregulated on granulomas formed by mycobacteria (53) which may promote resolution of inflammation was also observed in BCG-vaccinated Isg15+ macrophages. ISGs associated with immune clearance were also upregulated including *Snx10* (phagosomal maturation), *Gtgn2* (M1 polarization) and *Itgal* (T cell recruitment) were upregulated following BCG vaccination and BCG-vaccinated, SCV2-challenged hamsters (**Fig. 5D**). Lastly, Isg15+ macrophages from BCG-vaccinated, SCV2 challenged animals also showed up-regulation of genes associated with attenuation of inflammatory responses such as *Socs3*, *Il18bp*, *Dusp1* and *Fos* as well as those which may counter tissue damage phenotype induced by inflammation such as *Il1rn* (regulator of IL-1*α* and IL-1*β* cytokines) and *Timp1* (tissue inhibitor of MMP1) (**Fig. 5D**).

### DEGs in Siglecf+ AMs

In unvaccinated, SCV2-infected animals Siglecf*+* AMs highly expressed inflammatory genes (*Irf7*, *Irf9*, *Apobec1*, *Mx2*, *Mx1*, *Isg15*, *Ifit2*, *Rasd2*, *Herc6*, *Gbp5*, *Gbp7*, *Ifi44*, *Ifi47*, *Dhx58*, *Ifitm2*, *Igitp*, and *Cnp*) and genes involved in T cell recruitment and activation (*Ccl5*, *Ccrl2*, *Clec5a*, *Ltb*, *Ccl4CL4*, *Marcs1*, *Tnfsf10*, *Cd48*, *Alxox5*, *C3ar1*, *Cd80*, and *Cxcr2*). Genes characteristic of ER stress (*Gadd45*, *Gadd45b*, and *Atf4*) were also upregulated in *Siglecf*+ AMs of unvaccinated, SCV2-challenged animals (**Fig. 5E**). In the Siglecf+ AMs of these same animals we also noted expression of genes are associated with signaling to plasmacytoid DCs and monocytic DCs (*Unc93b1*, *Lsp1*, *Slamf9* and *Batf*) suggesting a program to keep these key sentinel cells in and activated state. In contrast, in Siglecf*+* AMs from BCG-vaccinated, SCV2-challenged hamster lungs we found upregulation of genes associated extracellular microbial sensing linked to NF*κ*B signaling (*Cox1*, *Cox2*, *Cox3*, *Tnfsf13b*, *Hsp90ab1*, *Apoe1*, *Gtgn2* and *Ckap4P*). In addition, genes involved in phagocytosis (*Mrc1*), antigen presentation (*H2aa*), and suppression of inflammation (*Cdo1*, *Mertk, Mrc1*) were also upregulated in *Siglecf*+ AMs from BCG-vaccinated, SCV2-challenged hamsters (**Fig. 5E**).

### Non-immune cells are modulated by BCG vaccination

Since we did not enrich for immune cells prior to scRNAseq analysis, we were able to investigate non-immune cell responses. We were able to identify 5 types of non-immune cells (AT2, endothelial, ciliated, AT1, and fibroblasts) in hamster lungs. We identified both type I alveolar cells (AT1 cells), which are non-replicative, thin flat squamous cells that form the basic structure of alveoli, as well as Type II cells (AT2 cells), which are cuboidal, release pulmonary surfactant, and may differentiate into AT1 cells during lung injury (54). Ciliated cells from bronchial epithelium, fibroblasts, and endothelial cells were also present.

Consistent with ongoing inflammatory lung injury during SCV2 infection at D4 and D7, we found that AT1, AT2, ciliated, and fibroblast cells were all significantly depleted in unvaccinated animals (**Fig. 6A**). Interestingly, BCG vaccination prevented this depletion suggesting a global protective effect by BCG against viral lung injury. Since this reversal of lung damage was most profound for AT2 cells, we investigated the gene ontology of these cells in BCG-vaccinated animals and found both an enhancement of genes involved in remodeling and response to injury as well as genes involved in immune responses (**Fig. 6B**). In contrast, among non-vaccinated animals the gene groups most strongly expressed in AT2 cells were those playing a role in viral responses and host defenses (**Fig. 6C**).

**Figure 6.**
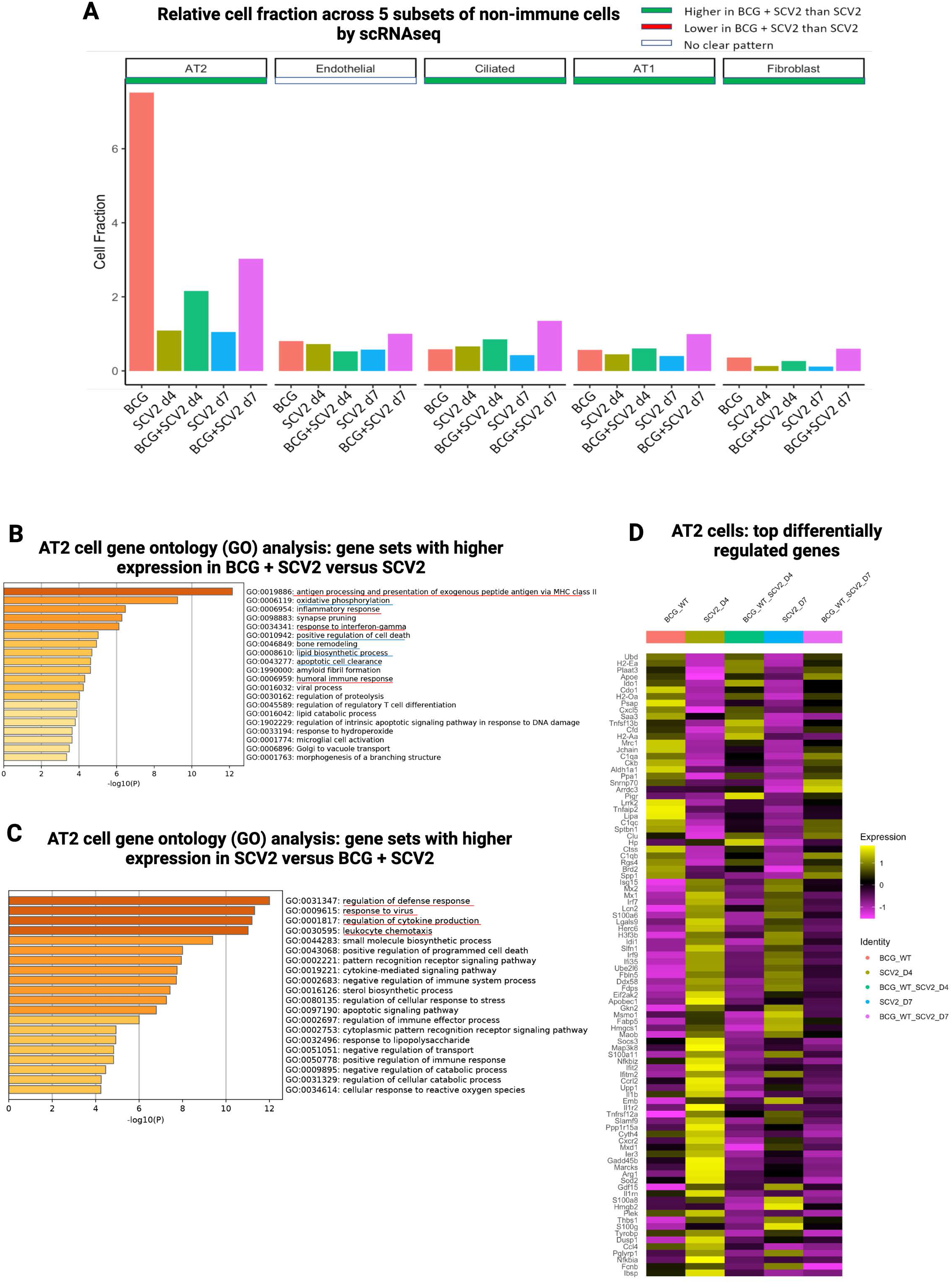
BCG vaccination restores AT2 cells and causes extensive phenotypic changes in non-immune cells in SARS-CoV-2 (SCV2) infected hamster lungs. **A**. Bar graph showing average proportion of different lung non-immune cells across groups. AT2 cell-linked gene ontology enrichment analysis of biological processes showing gene sets that enriched in (**B**) BCG + SARS-CoV-2 (SCV2) versus SARS-CoV-2 (SCV2), and in (**C**) SARS-CoV-2 (SCV2) versus BCG + SARS-CoV-2 (SCV2) across both time-points, and **D**. heat map depicting the average expression values of top differentially expressed genes (DEGs) across different groups in AT2 cells. Top DEGs were selected with adjusted P-value < 0.05 and average fold change > 0.7.

AT2 cells possess immunomodulatory functions and are considered the immune cells of the alveolar epithelium. Not surprisingly, when we evaluated DEGs in AT2 cells, we observed similar patterns to those seen in lymphoid and myeloid cells. In the absence of vaccination, prominent DEGs in AT2 cells were IFN-response genes (*Mx1*, *Mx2*, *Irf7*, *Irf9*, *Ifit2*, *Ifitm2*, *Slamf9*), antiviral response genes (*Apobec1*, *Ifit2,* and *Ddx58)* (**Fig. 6D**). In contrast, in BCG-vaccinated hamsters, AT2 DEGs were involved in immune clearance pathways such as antigen presentation (*H2-Ea*, *H2-Oa*, *H2Aa*, *Ctss* [cathepsin S, lysosomal Ag processing]), the complement pathway (*C1qa*, *C1qb C1qc*, *Cfd*), suppression of inflammation (*Ido1*), and lysosomal clearance (*Ubd*, *lipA* [lysosomal acid lipase]).

### Quantification of lung pathology using PET/CT

To evaluate pulmonary lung disease, SCV2 infected animals were imaged using ^18^F-FDG PET/CT at the peak of lung disease (D4) after infection. The ^18^F-FDG PET/CT analysis of lung consolidations did not reveal significant differences across the groups at D4 **(Fig. S7A-C)**.

## Discussion

In this study we evaluated the impact of intravenous BCG vaccination on the pathogenesis and immunology of SCV2 lung infection in the golden Syrian hamster model using viral quantification, histology, and flow cytometry supplemented by scRNAseq analysis. The SCV2 virus levels, histology and flow results showed that BCG vaccination prevented SCV2 replication at the peak of infection (D4) and that it also reduced the development of bronchopneumonia both at D4 and D7. Concomitantly, BCG vaccination blunted T cell lymphopenia in the lungs and reduced granulocyte lung infiltration. BCG vaccination was also associated with a significant recruitment of macrophages to the lung.

Our study is the first to use single cell transcriptional analysis in BCG-vaccinated, SCV2 infected animals. Our scRNAseq data revealed that BCG vaccination 4 weeks prior to SCV2 challenge was associated with significant shifts both in the populations of cell types present in the lungs and in the DEGs expressed by these cell types. Among lymphoid cells, we noted a unique lung recruitment of plasma cells in BCG vaccinated animals that was absent in SCV2-infected animals and also in BCG vaccinated animals that were not SCV2 infected. These plasma cells showed higher gene signature scores associated with elevated immunoglobulin production suggesting that they are involved in a BCG-mediated acceleration of antibody production against the SCV2. This expansion of plasma cells and humoral response may in part be an anamnestic reaction, since peptide epitopes from BCG proteins and SARS-CoV-2 NSP3 and NSP13 proteins have shown to share significant homology and this may allow for potential cross-reactive adaptive immunity (55, 56). Several categories of CD4 T cells (Th1, Th17, Treg, and Tmem cells) as well as CTLs were more abundant in the lungs of BCG vaccinated animals, and BCG-vaccination led to reduced exhaustion scores in the gene expression profiles of some of these T cells. Th1 and Tregs cells in the lung expressed high levels of type I IFN-associated genes in SCV2-infected animals as would be expected with viral infection; however, in these same cell types, among BCG-vaccinated, SCV2-infected lungs the predominant DEGs were shifted towards antigen presentation.

Among macrophages, scRNAseq revealed that BCG vaccination alone recruits high levels AMs to the lungs, a cell type associated with low inflammatory potential and clearance of pathogens. Upon SCV2 infection, BCG-vaccinated animals retained these high levels of AMs, while they dropped to low levels in SCV2-challenged, unvaccinated hamsters. In contrast we identified several populations of interstitial macrophages that were considerably more abundant in lungs of unvaccinated animals than in those that were BCG-vaccinated. As IM are non-resident macrophages likely recruited from the periphery which are known to have high inflammatory capacity, it is possible that a salutary immunologic effect of BCG is prevention of excess pro-inflammatory IM recruitment. While the DEGs of both the AM and IM populations in unvaccinated animals showed high expression of IFN-associated and chemokine genes, BCG vaccinated lungs showed AM and IM that strongly expressed metabolic and repair genes.

This study is the first to show that intravenous BCG protects against SCV2 infection in hamsters. We found a significant reduction of the SCV2 viral titer in BCG vaccinated animals at the peak of infection on day 4. A recent study of both intravenous and subcutaneous BCG in two species of hamsters (golden Syrian and Roborovski) found that neither route of vaccination protected either hamster species against SCV2 challenge (30). Our study differed from that of Kaufmann et al. in that we used a higher dose of BCG (5×10^6^ as opposed to 1×10^6^ CFU), a higher viral challenge dose (5×10^5^ as opposed to 1×10^5^ or 1.4×10^4^ PFU), and our peak disease sacrifice time was one day later (day 4) than theirs (day 3). Two mouse-based studies have evaluated BCG vaccination for SCV2 using either a mouse-adapted SCV2 virus or the K18-hACE2 mouse model. The first found no protection with subcutaneous BCG (28), while the second found that intravenous BCG but not subcutaneous BCG was protective (29). Despite being a well-defined animal model of COVID-19 with human-like pathology, one key limitation of using golden Syrian hamsters is the limited immunological resources such as well-characterized and validated antibodies (anti-CD8 Abs for example) and the relative paucity of immunologic data on hamster immune responses for correlation with our scRNAseq data. Along these lines, it was surprising that no CD8^+^ T cells were observed across the hamster groups even though they have been well-described in human scRNAseq studies of BAL samples in SCV2 (57, 58).

While BCG vaccination is routinely used in most countries as a TB prevention, the vaccine is generally given intradermally, and recent human studies showing protection by BCG against COVID-19 disease also used intradermal BCG (13). Like other animal studies using BCG in animal models of SCV2 (28) we used the intravenous route based on literature that IV BCG is far more potent against tuberculosis in non-human primates, and evidence that IV BCG reprograms bone marrow hematopoietic stem cells towards a more protective state against bacterial challenge (22). Nevertheless, SCV2 animal studies using percutaneous BCG have shown significant immunologic benefits at the level of flow cytometry (29), and we anticipate that many of the scRNAseq shifts which we observed in IV BCG vaccinated hamsters would be present following intradermal BCG albeit potentially at a lower level.

In summary, our study reveals that BCG vaccination reduces SCV2 replication and prevents bronchopneumonia in hamsters by mechanisms that involve enhanced numbers of lung alveolar macrophages, blunting of SCV2-mediated T cell lymphopenia, and reduced lung granulocyte infiltration. BCG appears to accelerate the appearance of immunoglobulin-producing plasma cells in the lung suggesting accelerated antiviral antibody production. The fact that BCG elevates the abundance of lung Treg cells while shifting a number of cell types away from expression of IFN-associated genes, suggests that BCG has immunotolerizing activity. These observations indicate that BCG vaccination may play a valuable role in protection against SCV2 and suggest that further studies of combining BCG with existing COVID-19 vaccines may offer synergistic protection.

## Methods

### Study approval

All experiments with infectious SCV2 were carried out in Institutional Biosafety Committee approved BSL3 and ABSL3 facilities at The Johns Hopkins University School of Medicine using recommended positive pressure air respirators and protective equipment. Experimental procedure involving live animals were carried out in agreement with the protocol (#HA20M310) approved by the Institutional Animal Care and Use Committee (IACUC) at The Johns Hopkins University School of Medicine.

### Bacterial strain and culture conditions

In this study we used commercially available *Mycobacterium bovis* (*M. bovis*) *Bacillus Calmette-Guérin* (BCG)-Tice (Onco-Tice^©^, Merck) for immunization experiments. The lyophilized bacterial stock was resuspended in 1ml of 7H9 Middlebrook liquid medium (Cat. B271310, Fisher Scientific) supplemented with (OADC) (Cat. B11886, Fisher Scientific), 0.5% glycerol (Cat. G55116, Sigma) and 0.05% Tween-80 (Cat. BP338, Fisher Scientific). The culture was streaked on 7H11 plate supplemented with oleic-albumin-dextrose-catalase (OADC) and single colonies were picked and propagated in 7H9 Middlebrook liquid medium for preparation of seed-stock. Individual seed-stock vial was randomly picked from frozen stock and was subsequently propagated in 7H9 medium before immunization.

### Cells and viruses

All cells were obtained from the American Type Culture Collection (ATCC, Manassas, VA, USA). Vero C1008 [Vero 76, clone E6, Vero E6] (ATCC CRL-1586) cells were used for viral growth and determination of virus stock titers. Vero E6 cells were grown in MEM with 10% fetal bovine serum (FBS), L-glutamine, and penicillin-streptomycin at 37°C with 5% CO_2_. SARS-CoV-2/Wuhan-1/2020 virus (U.S. Center for Disease Control and Prevention) was provided by Dr. Andrew Pekosz. The viral stocks were stored at -80°C and titers were determined by tissue culture infectious dose 50 (TCID_50_) assay.

### Animals

In vivo experiments involving BCG vaccination and SCV2 infection were carried out using male golden Syrian hamsters (*Mesocricetus auratus*). Male golden Syrian hamsters (age 5 to 6 weeks) were purchased from Envigo (Haslett, MI). Animals were housed individually under standard housing conditions (68 to 76°F, 30 to 70% relative humidity, 12-h light/12-h dark cycle) in cages with proper bedding (Teklad 7099 TEK-Fresh, Envigo, Indianapolis, IN) in the Animal Biosafety Level 3 (ABSL3) facility at Koch Cancer Research Building 2 (CRB2) at School of Medicine at The Johns Hopkins University. Animals were given ad libitum reverse osmosis (RO) water and feed (2018 SX Teklad, Envigo, Madison, WI). After 7 days of acclimatization animals were intravenously vaccinated using live 5 x 10^6^ C.F.U. of BCG-Tice in a total volume of 100 ul saline under ketamine (60 to 80 mg/kg) and xylazine (4 to 5 mg/kg) anesthesia administered intraperitoneally. Control animals received an equivalent volume of saline. Animals were challenged using 5 x 10^5^ TCID_50_ of SCV2/Wuhan-1/2020 virus in 100 μl of DMEM (50 μl/naris) through the intranasal route under ketamine (60 to 80 mg/kg) and xylazine (4 to 5 mg/kg) anesthesia administered intraperitoneally. Control animals received an equivalent amount of DMEM. Animals were randomly assigned to be euthanized by isoflurane overdose at end points (day 4 and day 7) following SCV2 infection.

### Experimental design

Groups of hamsters were intravenously (I.V.) administered saline or were vaccinated (I.V.) with 5×10^6^ CFU of BCG-Tice. 30 days after vaccination animals were challenged with 5×10^5^ TCID_50_ units of SARS-CoV-2 (SCV2) (Wuhan-1/2020) by the intranasal route. At D4 and D7 post-challenge animals were sacrificed for analysis. 3 animals (BCG only) were not infected with SCV2 and were sacrificed 30 days post-vaccination to serve as controls (**Fig. 1A**).

### Flow Cytometry analysis

For cellular immune profiling cell surface staining was performed on single cells from hamster lung and spleen tissues at the experimental end point. Briefly, tissues were harvested and stored in sterile PBS in individual tubes before single cell preparations. Lung was extensively perfused using sterile PBS. We used mouse lung (Miltenyi Biotec; 130-095-927) and spleen dissociation kits (Miltenyi Biotec; 130-095-926) for preparation of single cells as per manufacturer’s instruction using a gentleMACS™ Octo Dissociator with heaters (Miltenyi Biotec; 130-096-427). Cells were passed through a 70 μm filter and washed twice using ice-cold PBS followed by RBC lysis using ACK lysis buffer (Thermo Fisher Scientific: A1049201) at room temperature for 5 minutes. The cell viability was determined using Trypan blue dye staining and determine total live cells per lobe. For surface staining a total of 5 million cells per animal lung were used in this study. Briefly, cells were washed again using ice-cold PBS and stained using Zombie Aqua™ Fixable Viability Kit (Biolegend; 423101) for 20 min at room temperature. Cells were subsequently washed and resuspended in FACS buffer (1% BSA, 2mM EDTA in PBS) and incubated for 30 minutes at 4 °C in block buffer consisting of PBS and 2% FBS, 2% normal rat serum (Sigma Aldrich) and 2% normal mouse serum (Sigma Aldrich) prior to surface staining. Cells were again washed and stained with conjugated primary antibodies as per manufacturer’s protocol. Following antibodies were used for cell surface staining: anti-CD3 (Bio-Rad, #MCA1477PB, CD3-12), anti-CD4 (BioLegend, #100451, GK1.5), anti-CD11b (Novus, #NB110-89474PECY7, Poly), and anti-RT1D (BioLegend, #110211, 14-4-4S). Cells were subsequently fixed using IC fixation buffer (eBiosciences™; 00-8222-49) for 60 minutes at 4 °C. Cells were washed three times using FACS buffer and acquired using BD LSRII with FACSDiva Software. Data analyses was carried out using FlowJo (v10) (TreesStar).

### Sample and Preparation for scRNAseq

For single-cell RNA seq (scRNA-seq) of lung tissue derived single cells, whole right lung superior lobe was isolated from each animal. For single cell preparation, mouse lung dissociation kit (Miltenyi Biotec) was used with some additional modifications. Following single cell preparation, cell suspensions were applied to a MACS Smart Strainer (70 μm) and washed twice with 10 ml of DMEM. Cells were pelleted by spinning at 300 x g for a total of 7 minutes at 4°C. Cells were resuspended in 1ml DMEM containing 200 μl DNase (1μg/μl) for 5 minutes at room temperature. Cells were washed by adding 10 ml sterile PBS containing 0.5% BSA and pelleted at 4°C. The RBC lysis was carried out using 1X RBC lysis solution (Miltenyi Biotec; 130-094-183) in a total volume of 1ml for 10 minutes at 4°C. Cells were resuspended in 10 ml chilled DPBS containing 0.5% BSA and pelleted subsequently. Cells were filtered using 35 μm Falcon cell strainer and transferred to a 2 ml lo-binding tube in a total volume of 1ml DPBS containing 0.04% BSA. Cells were subsequently counted using trypan blue dye exclusion assay to determine cell viability and total cell number.

### Single-cell RNA sequencing and data pre-processing

Cells and gel beads were partitioned using the 10X Genomics Chromium platform aiming for recovery of 10,000 cells per sample using the Chromium Next GEM Single Cell 3ʹ Reagent Kits v3.1 (Dual Index) (CG000315). After RNA capturing, cDNA synthesis and library construction, the 3’DGE libraries were sequenced on NovaSeq 6000 instrument to achieve a target sequencing depth of ∼ 50,000 reads per cell.

Cell Ranger v4.0.0 was used to demultiplex the FASTQ reads and align them to the hamster transcriptome. The resulting gene expression matrix were then loaded into Seurat (v4.0.4) (59) in R (v4.0.3). ND1, ND2, ND4, ND5, ND6 were used to determine the percentage of mitochondrial genes. The quality of cells was assessed based on the number of genes detected per cell and the proportion of mitochondrial gene counts. Low-quality cells were filtered out if the number of detected genes were below 500 or above 5,000. Cells were filtered out if the proportion of mitochondrial gene counts was higher than 10%. In addition, genes that were expressed in less than 5 cells were excluded.

### Data integration and cell type identification

To compare cell types and proportions across different groups, we applied the integration methods using the SCTransform workflow, which can be used to assemble multiple distinct datasets into an integrated one and eliminate potential batch effect (60). Briefly, top 3000 highly variable features were identified and used to find anchors between each dataset with FindIntegrationAnchors function of Seurat. Then these anchors were used to integrate different datasets together with IntegrateData function to create a batch-corrected data assay for downstream analysis.

Principal component analysis (PCA) was performed on the integrated data for dimensionality reduction and top 30 significant PCs were used for cluster analysis. After using FindNeighbors and FindClusters functions, we performed nonlinear dimensional reduction with the RunUMAP function to obtain a two-dimensional representation of the cells. Cells with similar transcriptome were clustered together. The FindAllMarkers function in Seurat was used to find marker genes of each cluster for cell type identification. Clusters were then annotated based on the expression of canonical cell marker genes described in previous study (61) and a single cell sequencing database PanglaoDB (https://panglaodb.se/) (62). Basically, *Arg1*, *Ccr2* and *Ccr5* for MDSCs, *Camp* and *Mmp9* for granulocyte, *Siglecf*, *Marco* and *Mrc1* for alveolar macrophage (AM), *C1qb* for interstitial macrophages (IMs), *Cd3d* and *Cd4* for T cell, *Sftpb* and *Sftpc* for alveolar epithelial cell, *Nkg7* and *Klrd1* for NK cell, *Mki67* for proliferating cell, *Ms4a1* and *Cd79b* for B cell, *Cfap44* and *Foxj1* for ciliated cell, *Vcam1* and *Vwf* for endothelial cell, *Irf8* and *Tcf4* for dendritic cells (DCs), *Jchain* and *Mzb1* for plasma cell, *Dcn* and *Col1a1* for fibroblast. Clusters expressing two or more canonical marker genes characteristic of different cell types, were classified as doublets and excluded from further analysis. Lymphoid, myeloid, and non-immune cell subpopulations were further sub-setted and extracted, and separately analyzed by dimensionality reduction with PCA and UMAP.

To compare the abundance of each cell types across different samples, cell type proportion was calculated as the number of cells within each cell cluster divided by the total number of cells of that sample.

### Differentially expresses genes (DEGs) and pathway analysis

The FindMarker function in Seurat package was performed to identify differentially expressed genes between BCG-WT + SCV2 and SCV2 groups in a particular cell cluster. Benjamini–Hochberg method was used to adjust p-values for multiple tests. Genes expressed in at least 25% of cells in a cluster with adjusted P values < 0.05 were considered as differentially expressed genes. The gene filtering parameters used to generate each heatmap from differential gene expression analysis were described in each figure legend. GO term enrichment analysis of the DEGs was performed using Metascape (www.metascape.org) (63).

### UCell analysis

UCell (64) is an R package, based on the Mann-Whitney U statistic, for calculating gene signatures in single-cell datasets . We used UCell score to evaluate the degree to which individual cells expressed a certain gene set. HALLMARK_INTERFERON_ALPHA_RESPONSE from MsigDB, Immunoglobulin production involved in immunoglobulin-mediated immune response (GO:0002381) and 6 well-defined exhaustion associated genes (*Lag3*, *Tigit*, *Pdcd1*, *Ctla4*, *Havcr2* and *Tox*) (65) were used as gene signature input to calculate UCell scores for each individual cell. UCell score violin plots were grouped by different samples and cell types.

### Publicly available healthy control data

We additionally processed scRNA-seq data on hamster lung tissues by Nouailles et al. (61). Three datasets of lung samples from healthy hamsters (GSM4946629, GSM4946630, GSM4946631) were downloaded from the GEO database under accession code GSE162208. We processed the data using the similar strategy described above to filter low quality cells. Then MapQuery function in Seurat (v4.0.4) was performed to identify cell types using our own data reference. In brief, we transferred cell type labels from our reference data. Second, we integrated reference with query by correcting the low-dimensional embeddings. Finally, we projected the query data onto the UMAP structure of the reference, which allowed us to compare the abundance of the same cell type between datasets.

### Data availability

Our single cell sequencing data publicly available at NCBI through the GEO Series accession number GSE XXX [to be provided prior to publication]. The publicly available datasets that we used in this study are available from GEO GSE162208 (61).

### Histochemistry and Immunohistochemistry

Formalin-fixed paraffin embedded lung sections were stained with hematoxylin-eosin (H&E) for analysis. Lung tissue was scored using a panel of specific lung inflammation parameters according to criteria previous published on animal models of acute pneumonia (66). All examinations were performed by a board-certified pulmonary pathologist. Immunohistochemistry was performed on FFPE lung tissue sections for CD3 using an automated Ventana Discovery (Roche, Basel, Switzerland) autostainer. Heat induced antigen retrieval was achieved using ETDA pH9 solution (CC1, Roche). Primary antibody for CD3 (1:200 dilution; Cat# RM-9107-S1, Thermo Fisher Scientific) was incubated for 30m at 4 °C and detection was achieved using UltraView DAB detection kit (Roche). Protein expression was scored by a board-certified pulmonary pathologist blinded to cohort status or treatment group. CD3 membranous staining was evaluated separately in perivascular or peribronchial inflammation, bronchial epithelium, alveolar wall as well as pneumocytes and in areas of pneumonia (when present) or alveolar spaces (in absence of pneumonia). CD3 staining was scored as 1+ (scattered single cells), 2+ (cells in clusters or sheets with at least 2 layers) or 3+ (several clusters or sheets). Scores were then quantified using a point system (focal 1+= 0.5, 1+=1, focal 2+=1.5, 2+=2, 3+=3) and total points were analyzed between treatment groups. Total CD3 expression was calculated by combining all point scores for all criteria. Statistical analysis was performed between groups at each timepoint (day 4 post infection and day 7 post infection) using Welch’s t-test.

### SCV2 viral quantification

#### Quantification of SARS-CoV-2 infectious viral load in lung tissues

Following perfusion with phosphate-buffered saline (PBS), lung sections were collected, weighed and snap frozen on dry ice. Ice-cold virus titer buffer (Dulbecco’s Modified Eagle Medium, 2% fetal bovine serum, 2 mM L-glutamine, 200 U/mL penicillin and 200 μg/mL streptomycin, 100 μg/mL gentamicin, 0.5 µg/mL amphotericin B) was added to the frozen lung samples at a 10% weight to volume ratio. The tissue was homogenized with ceramic 1.4 mm beads for 2 cycles of 20 seconds at a speed setting of 5000 rpm using a Precelley’s Evolution homogenizer. Samples were centrifuged at 10,000 xg for 1 minute at 4°C to pellet the beads and tissue debris. In brief, lung homogenates were serially diluted in seven-point half-log dilutions and incubated on Vero C1008 cells in six technical replicates. Each sample was measured in triplicate. Plates were scored for CPE five days post infection using the CellTiter-Glo® Luminescent Cell Viability Assay per the manufacture’s protocol (Promega). TCID_50_ was calculated using the Reed and Muench method (67).

#### RNA extraction and RT-qPCR

Total RNA from hamster lung homogenates were extracted using TRIzol Reagent (Invitrogen) followed by the RNeasy kit (Qiagen) according to the manufacturer’s protocol. cDNA was synthesized from total RNA using qScript cDNA SuperMix containing random hexamers and oligo-dT primers (Quanta Biosciences) following the manufacturer’s protocol. Real-time PCR was performed in triplicate using TaqMan Fast Advanced Master Mix (Applied Biosystems) on a StepOnePlus Real Time PCR system (Applied Biosystems). SARS-CoV-2 RNA was detected using premixed forward (5’-TTACAAACATTGGCCGCAAA-3’) and reverse (5’-GCGCGACATTCCGAAGAA-3’) primers and probe (5’-FAM-ACAATTTGCCCCCAGCGCTTCAG-BHQ1-3’) designed by the CDC as part of the 2019-nCoV CDC Research Use Only (RUO) kit (Integrated DNA Technologies, Catalog #10006713) to amplify a region of the SARS-CoV-2 (SCV2) nucleocapsid (N) gene. PCR conditions were as follows: 50°C for 2 min, 95°C for 2 min, followed by 45 cycles of 95°C for 3 s and 55°C for 30 s. Serially diluted (10-fold) plasmid containing the complete SARS-CoV-2 N gene (Integrated DNA Technologies, Catalog #10006625) was measured to generate a standard curve for quantification of viral RNA copies. The limit of detection for the assay was 1 x 10^2^ RNA copies. Viral copies were normalized to the human RNase P (RP) gene using premixed forward (5’-AGATTTGGACCTGCGAGCG-3’) and reverse (5’-GAGCGGCTGTCTCCACAAGT-3’) primers and probe (5’-FAM-TTCTGACCTGAAGGCTCTGCGCG-BHQ-1-3’) included in the same 2019-nCoV CDC RUO kit.

#### CT and PET imaging

Four days post infection, live SCV2-infected male hamsters (n= 5 SCV2 and n= 6 BCG + SCV2) underwent chest CT using the nanoScan positron emission tomography (PET)/CT (Mediso USA, MA, USA) small animal imager. Prior to PET imaging, hamsters were administered ∼10.22 MBq of ^18^F-FDG (SOFIE, Sterling VA, USA) via the surgically implanted center venous catheter. Given that SCV2 is designated as a BSL-3 pathogen, live SCV2-infected animals were imaged inside transparent and sealed biocontainment cells developed in-house, compliant with BSL-3 containment and capable of delivering air-anesthetic mixture to sustain live animals during imaging (68, 69). A 15-min PET acquisition and subsequent CT were performed using the nanoScan PET/CT (Mediso Alrington, VA). CT images were visualized and analyzed using the VivoQuant 2020 lung segmentation tool (Invicro, MA, USA) (69). Briefly, an entire lung volume (LV) was created, and volumes of interests (VOIs) were shaped around the pulmonary lesions using global thresholding for Hounsfield Units (HU) ≥ 0, and disease severity (CT score) was quantified as the percentage of diseased lung in each animal. The investigators were blinded to the group assignments. The data are represented as CT score [(pulmonary lesions volume/whole lung volume) × 100]. The investigators analyzing the CT were blinded to the group assignments. VivoQuant™ 2020 (Invicro, Boston, MA, USA) was used for visualization and quantification. Scatter and attenuation corrections were applied to the PET data and multiple VOIs were manually drawn per animal using the CT as a reference.

## Author contributions

A.K.S., R.W, W.R.B., S.Y., and T.J.B co-led the study through conceptualization, design, oversight, and interpretation of results. W.R.B. obtained the funding for the study. A.K.S. designed, conducted, and interpreted the results of the experiments. A.K.S., R.W., K.A.L., M.P. C.K.B, P.U., A.A.O., S.D., O.K., M. B., P.I., and K.J.P. conducted then experiments and provided key experts advice. S.K.J., and T.J.B. assisted in the design of the experiments and provided key expert advice. A.K.S., and W.R.B wrote the manuscript. A.K.S., R.W. K.A.L., M.P., C.K.B., O.K., A.A.O., K.J.P., S.K.J., T.J.B., S.Y and W.R.B. revised and edited the manuscript. A.K.S., and W.R.B. designed and produced figures for this manuscript.

## Acknowledgements

The generous support of grants AI155346 from the NIH and grant #2167 from Emergent Ventures at the Mercatus Center, George Mason University is gratefully acknowledged. This study was also supported by NIH grant U54CA260492 (providing support to S.Y. and R.W.). We thank the members of the Sidney Kimmel Comprehensive Cancer Center’s Experimental and Computational Genomics Core, supported by NIH Grant P30CA006973, for their support of the next generation sequencing experiments. The authors also thank Geetha Srikrishna, Ada Tam and Lee Blosser for technical assistance.

---

**Supplementary Figure S1.**
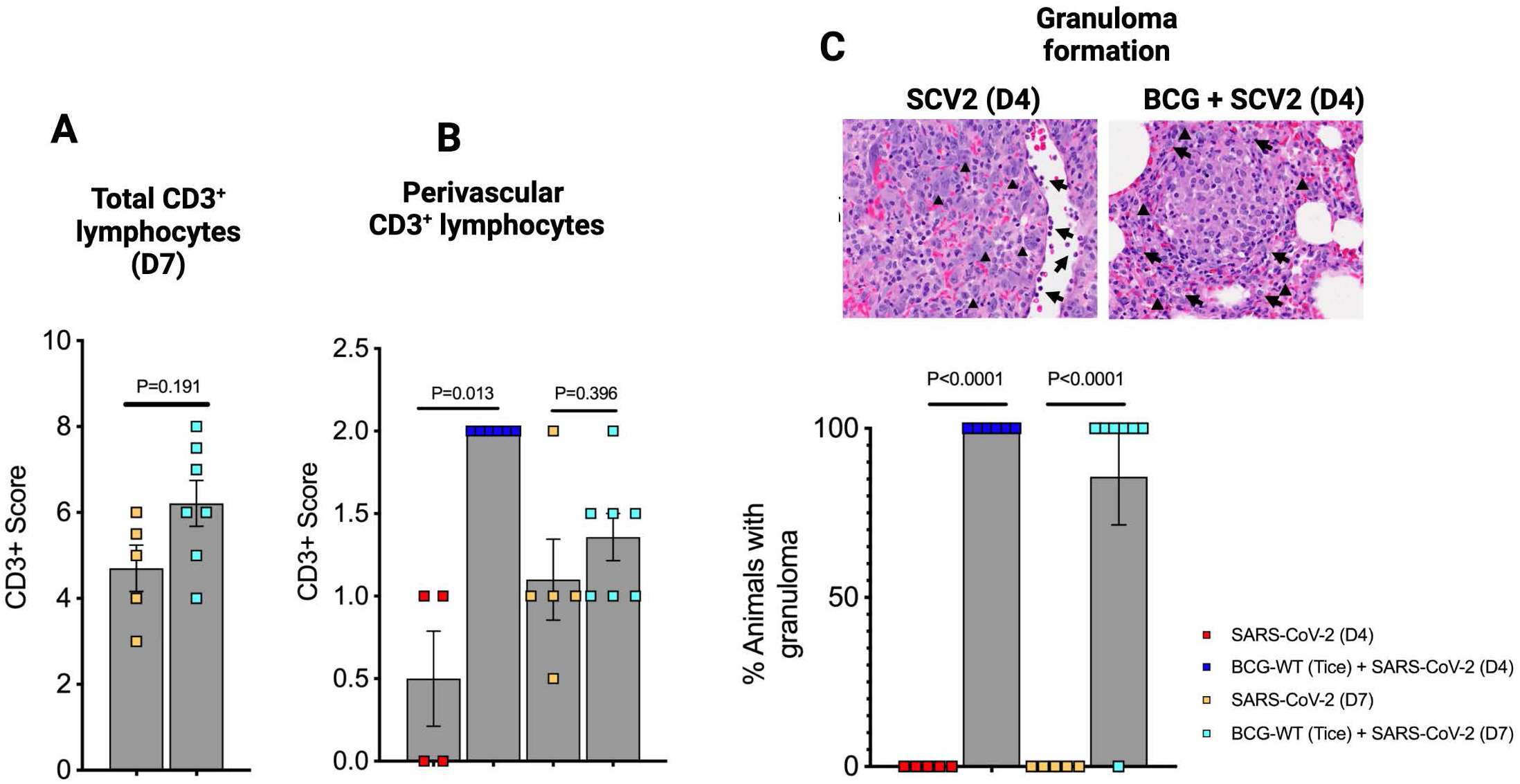
Lymphocyte infiltration and granuloma formation in BCG vaccinated golden Syrian hamsters after SARS-CoV-2 infection. **A.** total CD3^+^ lymphocytes at day 7 (D7). **B.** CD3^+^ lymphocytes in perivascular area, and **C.** Granuloma formation and inflammation. Left panel showing HPF (200x) view of SARS-CoV-2 infected lungs showing alveolated parenchyma with marked pneumocyte atypia (arrowheads) acute inflammation (neutrophils showed by arrows) and breakdown of architecture sonsistent with acute lung injury secondary to pneumonia. Right panel HPF showing intraalveolar small non-necrotizing granuloma surrounded by abundant lymphocytes (arrowheads) and macrophages (arrows) in hamster lung at day 4 after SARS-CoV-2 challenge. Bar graph showing pathological quantification of granulomata after BCG vaccination. Data points represent number of animals per grouop. Data presented as mean values ± S.E.M. Statistical analyses done using Welch’s t-test (P value < 0.05 considered significance). Schematic of CD3 positivity is given in methods section.

**Supplementary Figure S2.**
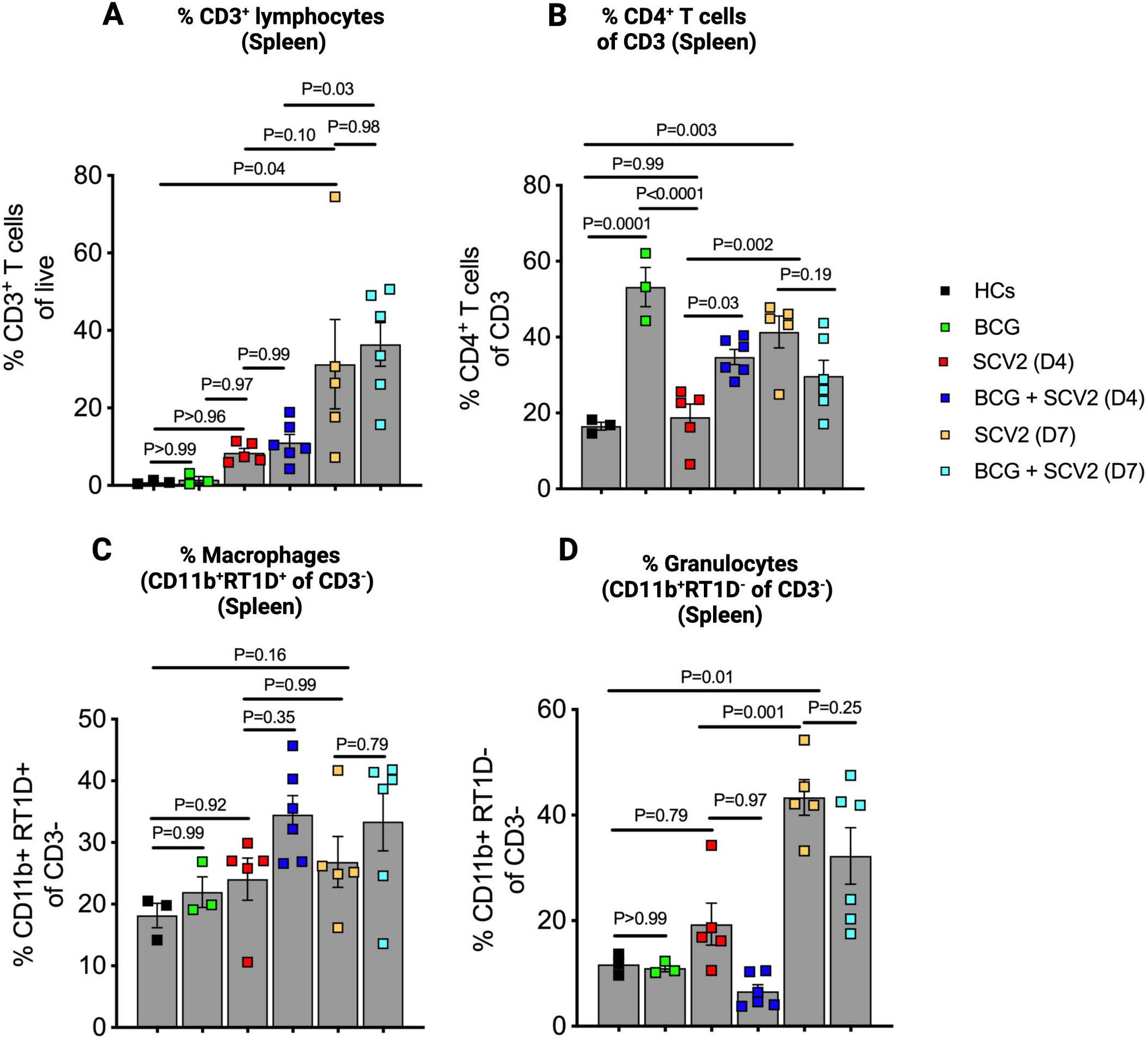
Differential abundance of lymphocytes and myeloid cells in hamster spleens across different groups. Bar graph showing percentages of **A.** total lymphocytes (CD3^+^). **B.** CD4+ T cells within lymphocytic compartment (CD4^+^ of CD3). **C.** macrophages (CD11b^+^RT1D^+^ of CD3^-^), and **D.** granulocytes (CD11b^+^RT1D^-^ of CD3^-^) within non-lymphocytic compartment. Data points represent the number of animals assigned per group. Data are represented as mean values ± S.E.M. The statistical analyses were done using one-way ANOVA (P values <0.05 was considered significant).

**Supplementary Figure S3.**
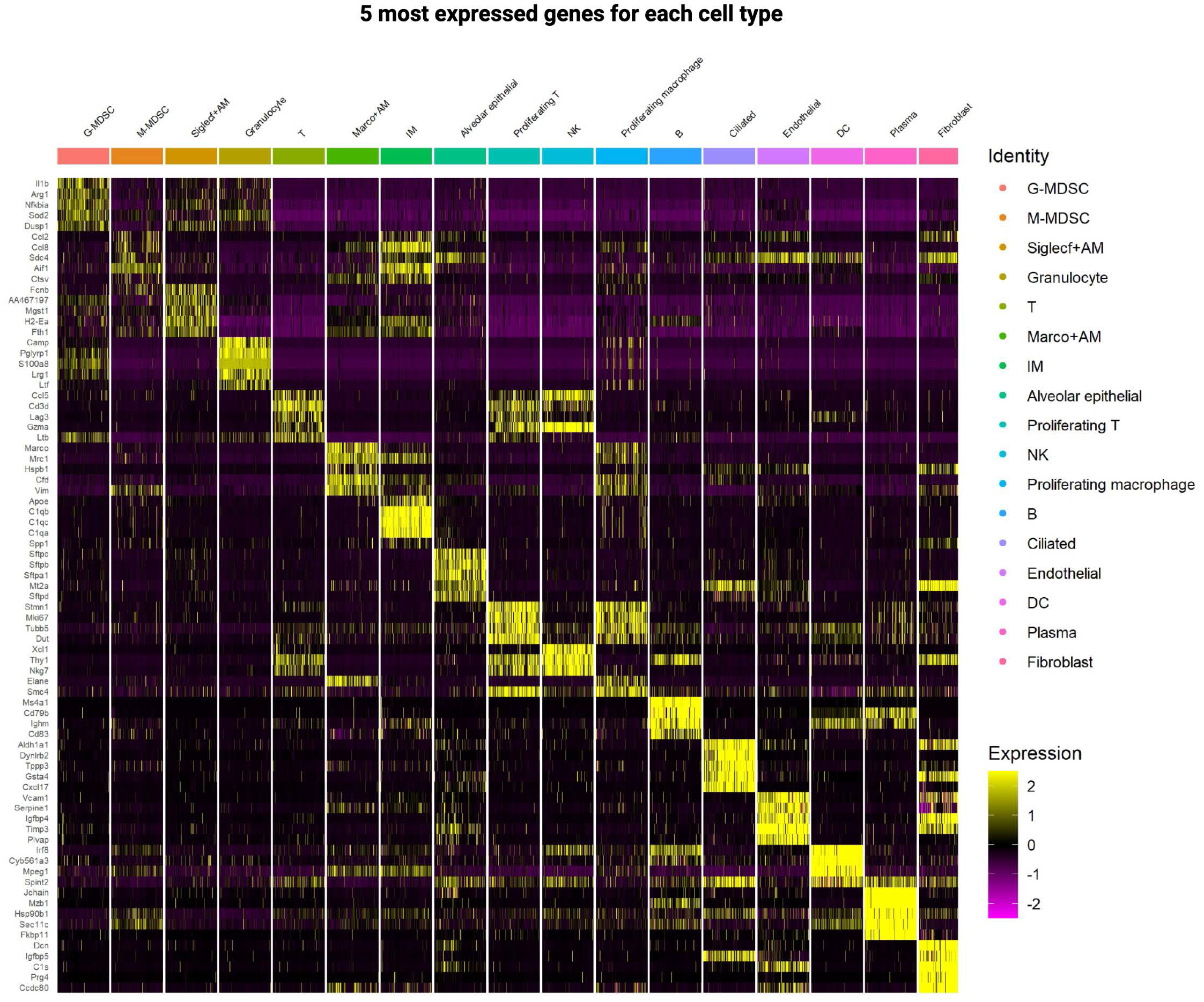
Heat map showing top 5 most upregulated genes across 17 cell types identified within immune (myeloid and lymphoid) and non-immune cells using single cell RNA sequencing (scRNAseq) analysis.

**Supplementary Figure S4.**
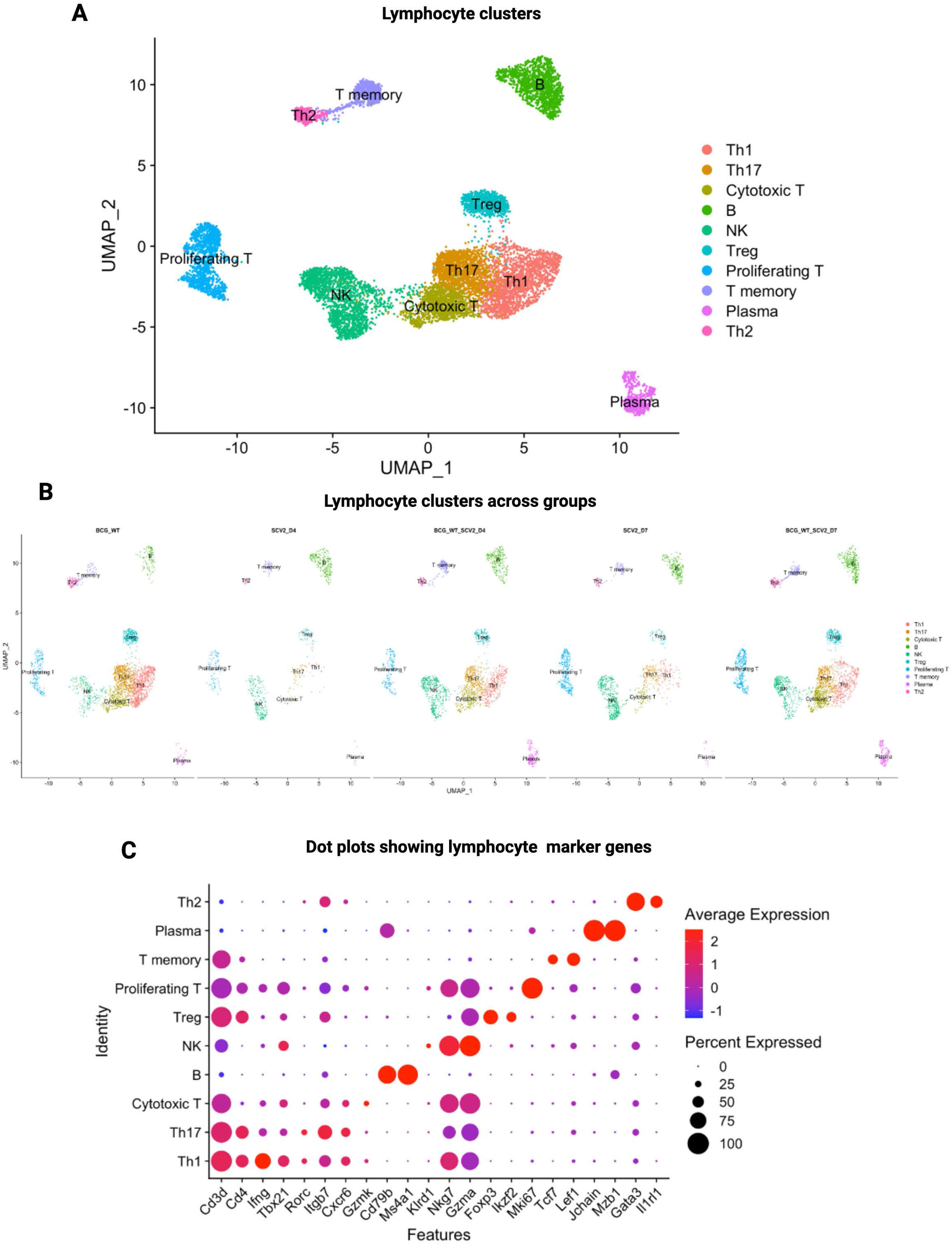
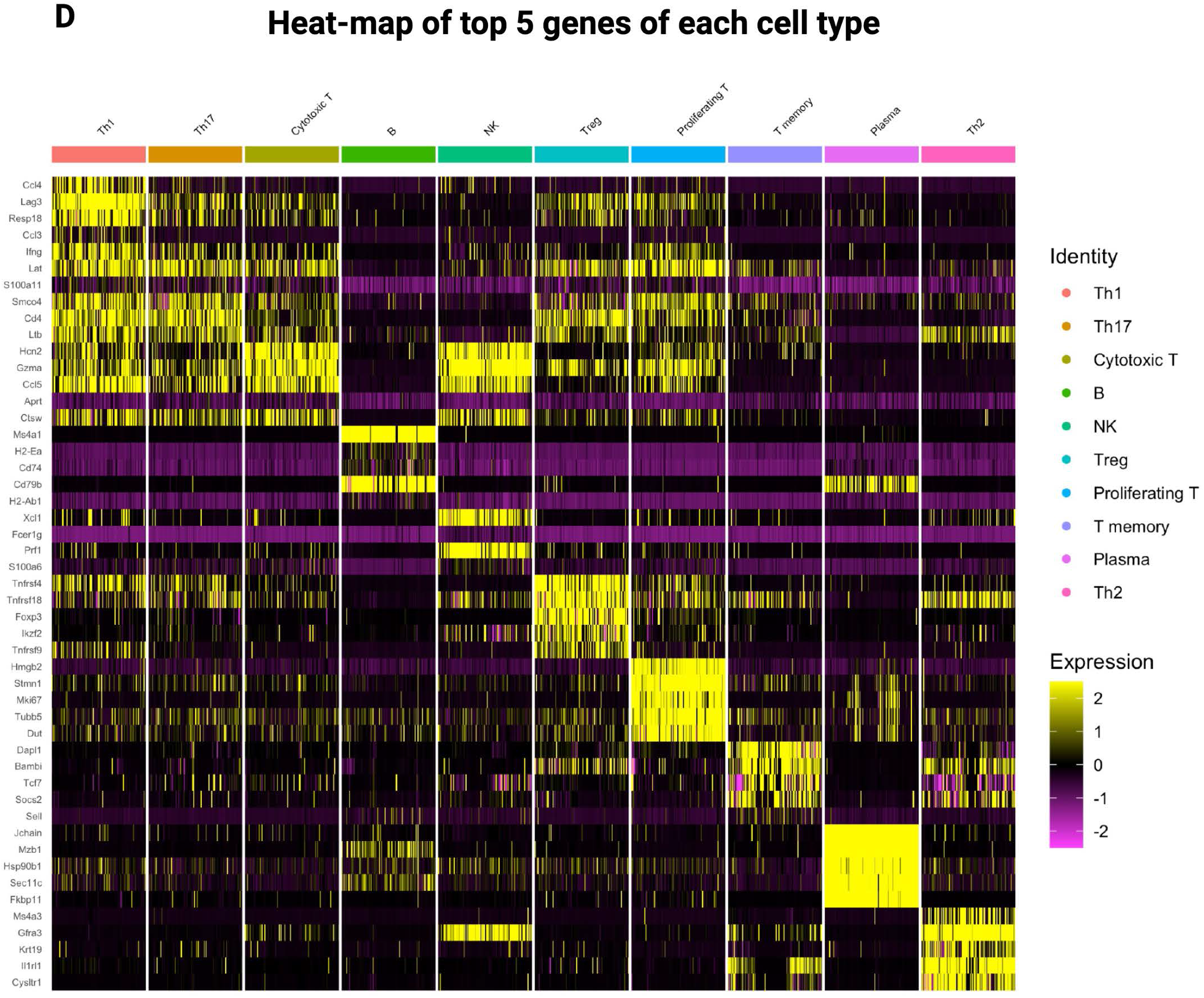
Identification of different lymphocytic sub-set of cells across groups. **A.** Uniform Manifold Approximation and Projection (UMAP) plot showing 10 lymphocytic sub-sets using the cognate markers. **B.** UMAP plot showing dynamic changes in abundance of different lymphocytic cells across groups at both time points. **C.** Dot plot showing average expression of markers genes used to assign cellular identity, and **D.** Heat map showing top 5 genes expressed in different lymphoid subsets.

**Supplementary Figure S5.**
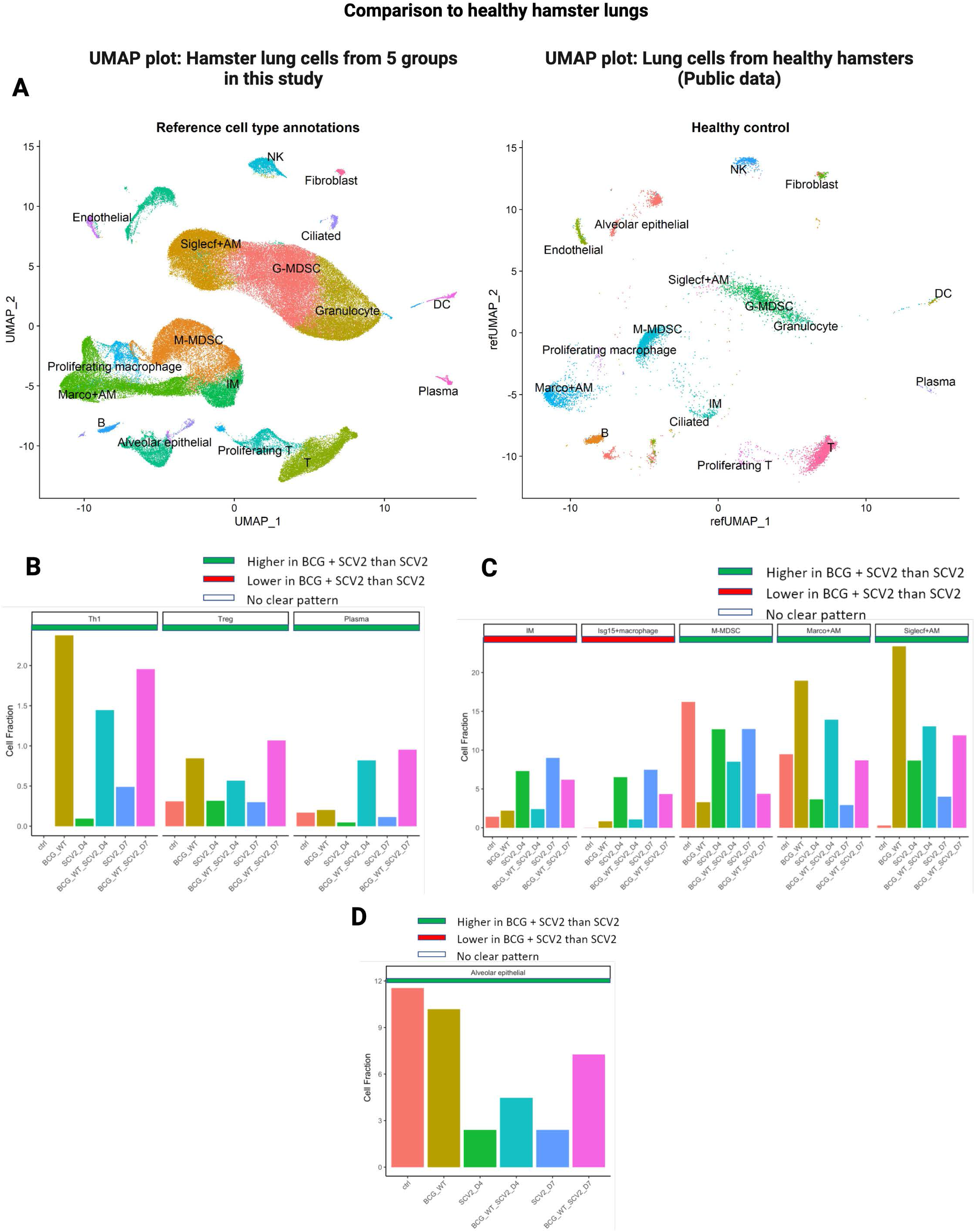
BCG vaccination induced remodeling of myeloid cells in animal lungs after SARS-CoV-2 infection. **A.** Uniform Manifold Approximation and Projection (UMAP) plot showing different myeloid cells in hamster lungs after SCV2 infection (this study) and in healthy age-matched hamsters (without SCV2 infection, public database). Average proportion of **B.** Th1, Treg and plasma cells. **C.** Major myeloid subsets, and **D.** Alveolar epithelial cells in hamster lungs across different groups.

**Supplementary Figure S6.**
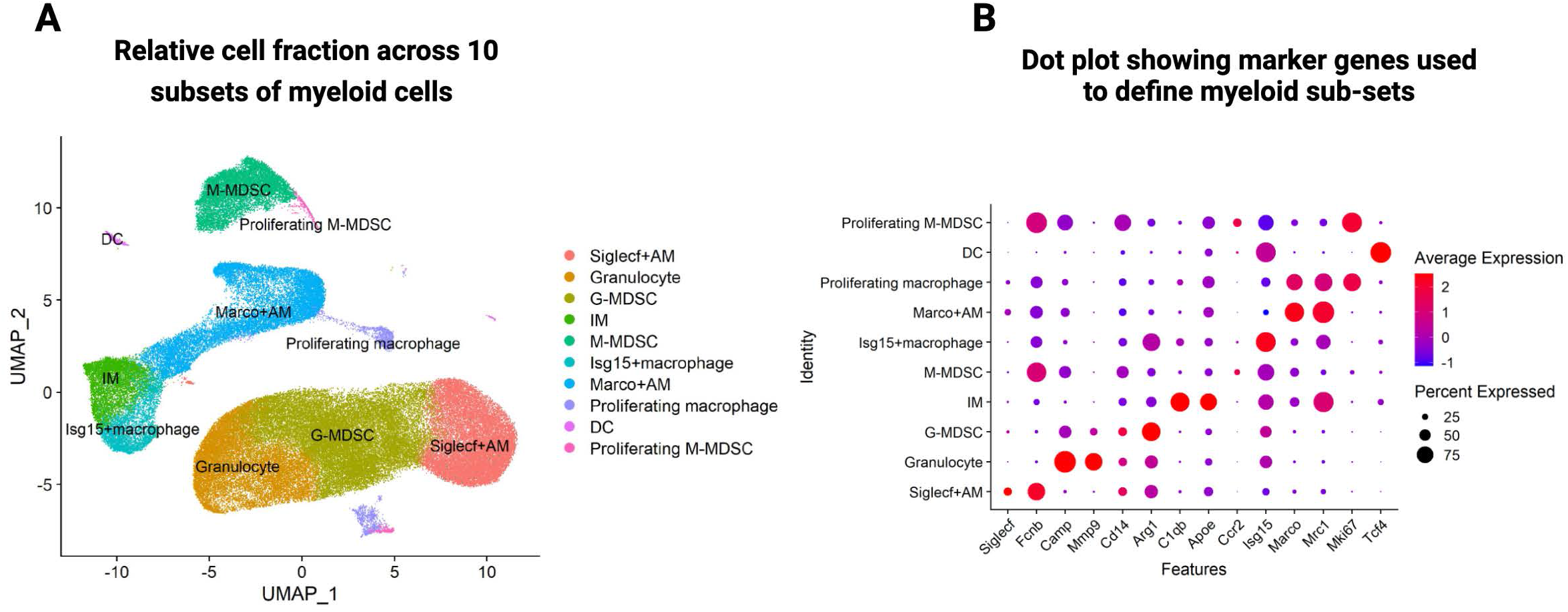
Identification of different myeloid sub-sets of cells across all samples. **A.** Uniform Manifold Approximation and Projection (UMAP) plot showing different myeloid subset of cells integrated across all samples. **B.** Dot plot showing average expression and percentage cells expressing marker genes used to define different myeloid sub-sets.

**Supplementary Figure S7.**
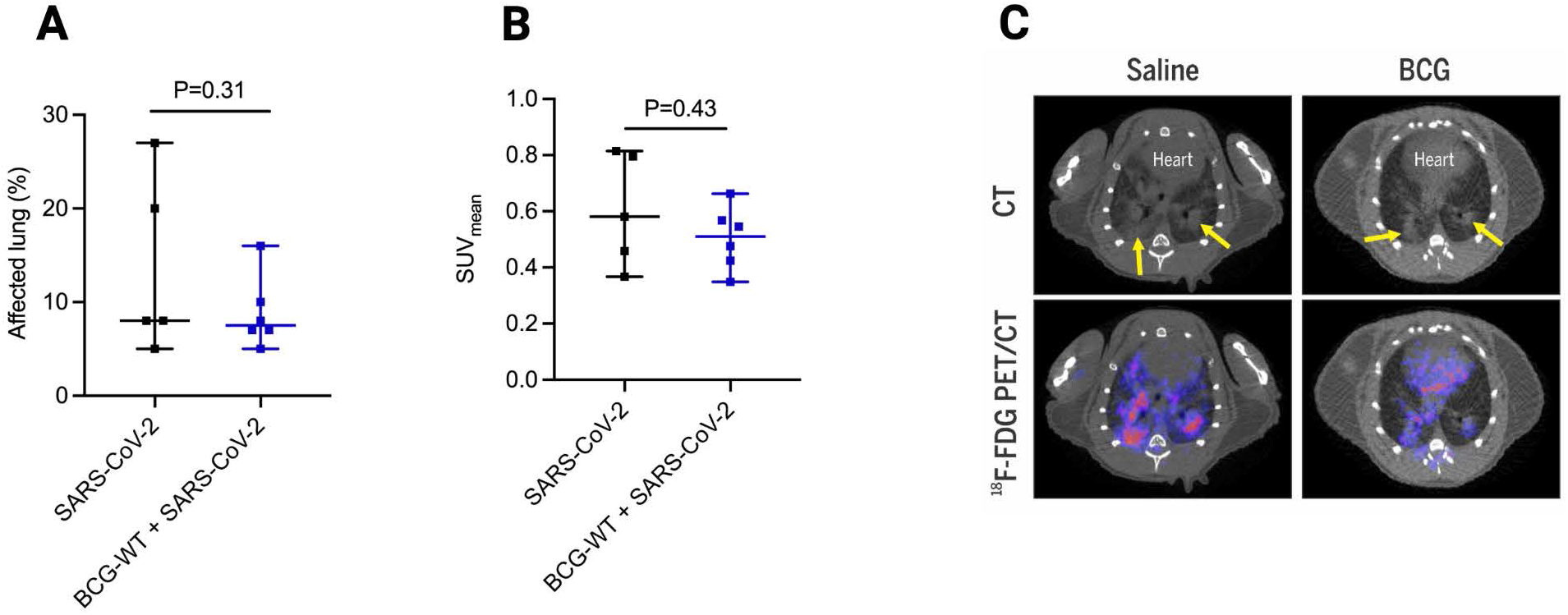
Monitoring lung pneumonia in hamsters infected with SARS-CoV-2 using ^18^F-FDG PET/CT. **(A)** The affected lung areas were determined for each SARS-CoV-2-infected hamster using CT. Automated lung segmentation of the CT images was performed and data is represented as percentage of the lung determined to be affected based on tissue density. **(B)** ^18^F-FDG PET/CT was performed in SARS-CoV-2-infected hamsters. The ^18^F-FDG PET signal in affected lung areas, as determined by the automated lung segmentation, was quantified, and is represented as mean standardized uptake value (SUVmean). The data is represented as median ± interquartile range. **(C)** Transverse ^18^F-FDG PET/CT of representative SARS-CoV-2-hamsters from the BCG-vaccinated and control groups. Increased ^18^F-FDG PET signal was observed in the areas of lung consolidation observed in CT (yellow arrows).

**Table S1:**
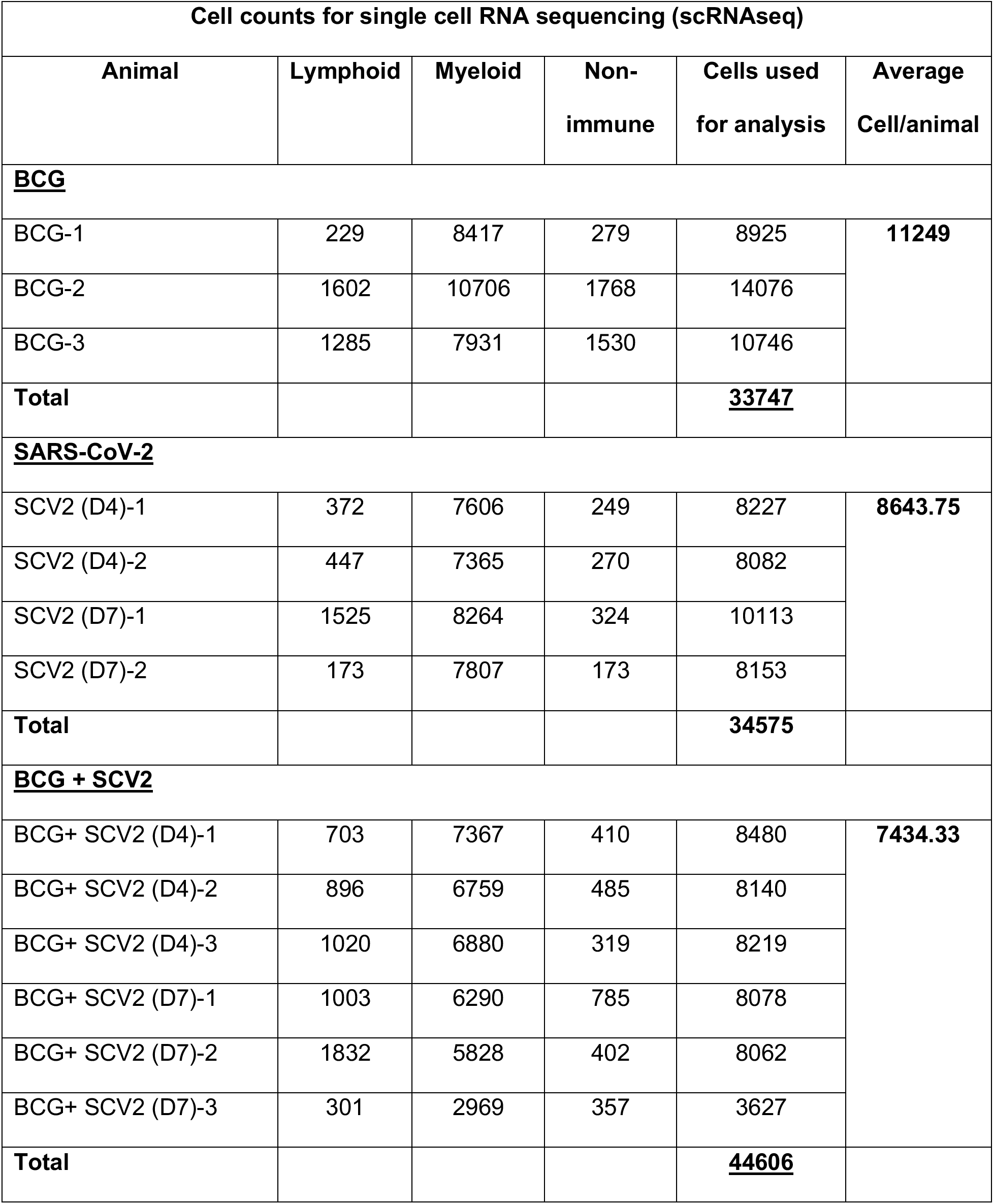
Table showing cell counts each in different immune and non-immune subsets

**Table S2:** Table showing identified cell types in this study based on specific marker genes

| Identification of different cell types based on marker genes |  |  |
| --- | --- | --- |
| No. | Cell Type | Marker Genes |
| **Immune cells (Myeloid)** |  |  |
| 1 | Granulocyte | *Mmp9, Camp* |
| 2 | G-MDSC | *Cd14, Arg1* |
| 3 | Marco+AM | *Marco, Mrc1* |
| 4 | IM | *C1qa, C1qb* |
| 5 | M-MDSC | *Ccr2, Ccr5* |
| 6 | Siglecf+AM | *Siglecf* |
| 7 | Dendritic Cells | *Tcf4* |
| 8 | Proliferating macrophages | *Macrco, Mki67* |
| **Immune cells (Lymphoid)** |  |  |
| 8 | T cells | *Cd3d, Cd4* |
| 9 | NK | *Nkg7, Prf1* |
| 10 | <u>B cells</u> | *Cd79b, Ms4a1* |
| 11 | <u>Plasma cells</u> | *Jchain, Mzb1* |
| **Non-immune cells** |  |  |
| 12 | Alveolar epithelial | *Sftpb, Sftpc* |
| 13 | <u>Ciliated</u> | *Foxj1, Cfap44* |
| 14 | <u>Endothelial</u> | *Vcam1, Vwf* |
| 15 | <u>Fibroblast</u> | *Dcn, Col1a1* |
| 17 | <u>Proliferating cells</u> | *Mki67, Top2a* |

